# Cover crop root channels promote bacterial adaptation to drought in the maize rhizosphere

**DOI:** 10.1101/2025.05.05.651044

**Authors:** Debjyoti Ghosh, Yijie Shi, Iris M. Zimmermann, Katja Holzhauser, Martin von Bergen, Anne-Kristin Kaster, Sandra Spielvogel, Michaela A. Dippold, Jochen A. Müller, Nico Jehmlich

**Affiliations:** Department of Molecular Toxicology, Helmholtz Centre for Environmental Research – UFZ GmbH, Permoserstraße 15, 04318 Leipzig, Saxony, Germany; Institute of Plant Nutrition and Soil Science, Department of Soil Science, Christian-Albrechts-University Kiel, Hermann-Rodewald-Straße 2, 24118 Kiel, Schleswig-Holstein, Germany; Institute of Crop Science and Plant Breeding, Agronomy and Crop Science, Christian-Albrechts-University Kiel, Am Botanischen Garten 1-9, 24118 Kiel, Schleswig-Holstein, Germany; Institute for Biochemistry, Faculty of Biosciences, Pharmacy and Psychology, University of Leipzig, Brüderstraße 34, 04103 Leipzig, Saxony, Germany; German Centre for Integrative Biodiversity Research (iDiv) Halle-Jena-Leipzig, Puschstraße 4, 04103 Leipzig, Saxony, Germany; Institute for Biological Interfaces, Karlsruhe Institute of Technology, Hermann-von-Helmholtz Platz 1, 76344 Eggenstein-Leopoldshafen, Baden-Württemberg, Germany; Geo-Biosphere Interactions, Department of Geosciences, University of Tübingen, Schnarrenbergstraße 94-96, 72076 Tübingen, Baden-Württemberg, Germany

**Keywords:** Cover crop, root channel re-use, bacterial community, drought, soil types, metaproteomics

## Abstract

**Background:** Increasing drought frequency poses a significant threat to agricultural productivity. A promising strategy to enhance crop resilience against drought is the utilisation of root channels left by winter cover crops, which can improve access to subsoil water and nutrients for subsequent cash crops like maize (*Zea mays* L.). The impact of drought on bacterial communities inhabiting these root channels remains largely unknown. Here, we investigated drought-induced shifts in maize rhizosphere bacterial communities and their functional adaptation in cover crop root channels across three soil types in northern Germany (Luvisol, Podzol, and Phaeozem).

**Results:** Using a multi-omics approach (16S rRNA gene amplicon sequencing, qPCR, and metaproteomics), we identified significant taxonomic and functional responses to drought. A rise in the abundance of *K*-strategist bacterial communities indicate a shift towards stress-tolerant populations with drought. Under drought stress, the relative abundances of *Acidobacteriota*, *Actinomycetota*, *Planctomycetota*, and *Pseudomonadota* increased, while *Chloroflexota*, *Methylomirabilota*, *Ca.* Patescibacteria, and *Verrucomicrobiota* declined. Functional analyses revealed that drought-stressed aerobic taxa among the *Pseudomonadota* and *Verrucomicrobiota* upregulated the glyoxylate cycle, potentially enhancing carbon and energy conservation, and increased antioxidant defences (catalase–glutathione peroxidase and methionine cycle–transsulfuration pathway). These drought-mitigating strategies were especially pronounced in root channels formed by *Brassicaceae* and *Poaceae* cover crops in the Luvisol and Podzol soils.

**Conclusions:** These findings demonstrate the functional plasticity of rhizosphere bacterial communities in reused root channels in response to drought, highlighting the potential to leverage microbiome-mediated resilience for agricultural practices.

**Graphical Abstract:** 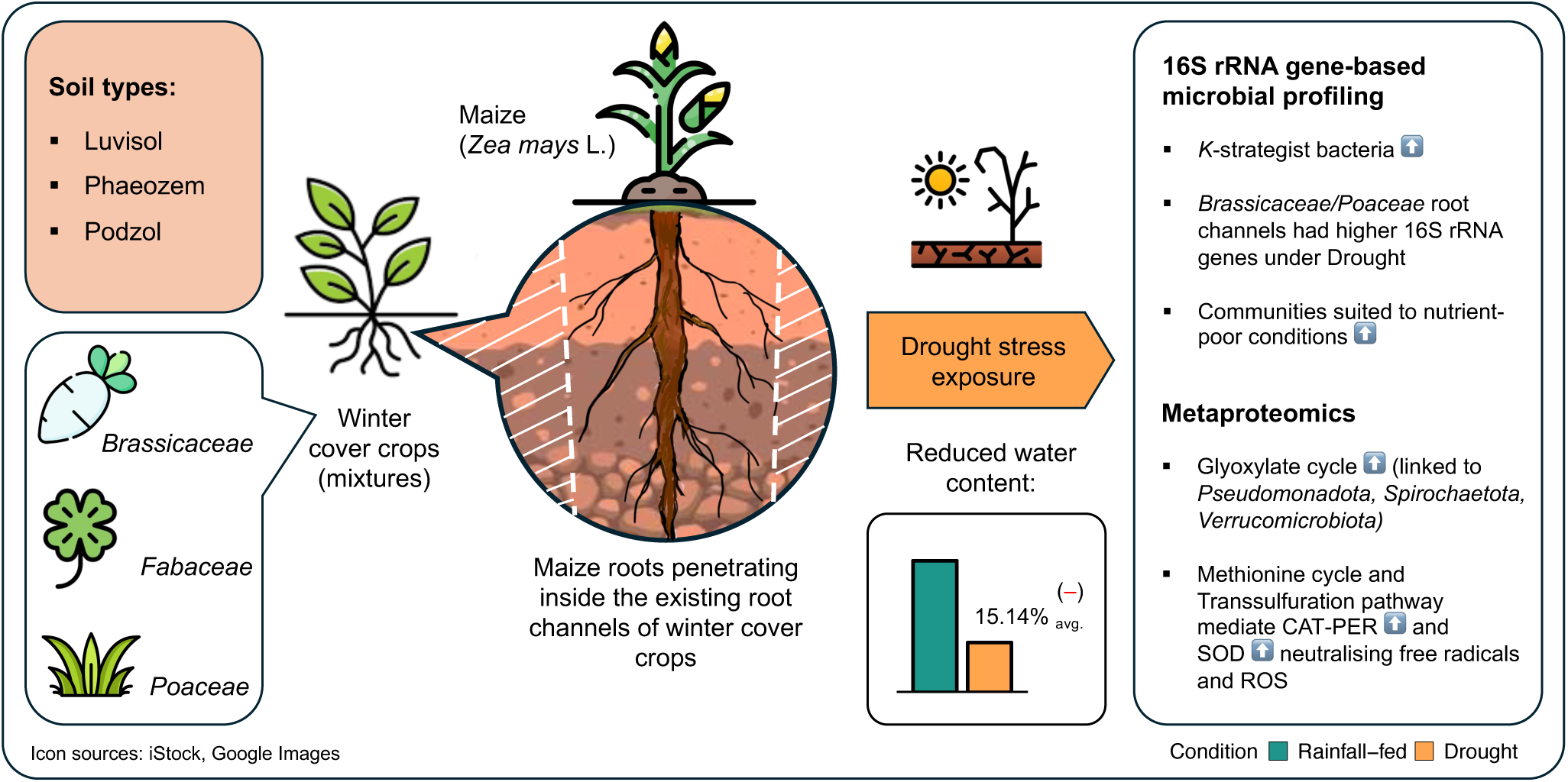

## Introduction

Increasing drought frequency and intensity, driven by climate change, significantly impact crop production worldwide [1-3]. Drought primarily impacts topsoil [4, 5], reducing water availability and limiting nutrient mobility and uptake [6]. Enhancing plant’s access to subsoil resources offers a promising strategy to mitigate these effects and sustain crop yields [7, 8]. Utilizing root channels created by previous cover crops can facilitate deeper root penetration and subsoil resource acquisition by subsequent cash crops [9]. These channels are recognized hotspots of microbial activity, exhibiting enhanced biogeochemical cycling compared to bulk soil [10]. In channels with decaying roots, the microbial activities are largely driven by residual complex root exudates and rhizodeposits, while in channels with viable roots, new labile plant-derived organic carbon mainly serves as microbial substrate [11]. Upon using maize a as model cash crop, we showed previously that bacterial abundances and functional diversity increase in the rhizosphere when the maize root when utilizing cover crop root channels compared to plants grown in unconditioned bulk soil [9].

Soil texture (loamy, clayey, sandy) significantly influences crop cultivation and yield by affecting root growth, water and nutrient uptake, and microbial community structure [12-14]. Under drought conditions, the bacterial community structure may shift in response to altered resource availability affected by texture-dependent drying out of the soil [15, 16]. Whether the structure, function, and dynamics of bacterial communities inhabiting root channels are similarly influenced by soil texture, or whether the channel properties themselves are predominant, remains poorly understood, particularly under water stress. Understanding these interactions is crucial for optimizing cover cropping strategies and enhancing crop resilience to drought.

To address this knowledge gap, we conducted field experiments using maize as a cash crop following mixtures of winter-grown cover crops. Drought conditions were simulated in a rainout shelter experiment. We employed 16S rRNA gene amplicon sequencing, quantitative PCR (qPCR), and metaproteomics to examine the effects of drought (D) compared to rainfall-fed (RF) maize cultivation on the structure and function of the rhizosphere bacterial community in maize reusing pre-existing root channels across different soil textures. We hypothesised that the bacterial microbiome of the maize rhizosphere reusing cover crop root channels is more activity and better in implementing adaptive mechanisms to mitigate drought-induced stress in a soil-type specific manner. This research aims to elucidate rhizosphere microbiome dynamics and adaptations in the biochemical pathways in topsoil and subsoil with varying textures under drought conditions, ultimately informing strategies for selecting cover crops and enhancing crop resilience in water-limited environments.

## Results

### Soil types, water content, and bacterial community diversity

The field study was conducted on sites with three different soil types, Luvisol, Phaeozem, and Podzol, in Germany in 2023. Plots were planted with cover crop mixtures with *Brassicaceae*, *Fabaceae* or *Poaceae* or left fallow, followed by cultivation of maize. Two soil profile depths, topsoil (0-30 cm) and subsoil (30-90 cm) were sampled during R1-RX growth stage of maize (bolting) (Fig. 1). The organic C content in the samples varied significantly with soil type and depth (Fig. S1, Table S1). Top- and subsoil total organic C (TOC) were highest in the Podzol followed by Luvisol and Phaeozem (topsoil: 1.76 _Podzol_ > 1.27 _Luvisol_ > 1.23 _Phaeozem_, subsoil: 0.95 _Podzol_ > 0.82 _Luvisol_ > 0.59 _Phaeozem_, units in g C kg^-1^soi). Total nitrogen (TN) content in the topsoil was the same across all three sites (0.13 g N kg^-1^ soil), but was higher in Luvisol subsoil than in the other two subsoil types (0.08 _Podzol_ > 0.06 _Luvisol_ ≥ 0.06 _Phaeozem_, units in g N kg^-1^ soil). Soil pH varied considerably, with Podzol being the most acidic in topsoil and subsoil (both pH 5.21), followed by Luvisol (topsoil pH 6.49, subsoil pH 6.48), and Phaeozem being neutral to slightly basic (topsoil pH 7.14, subsoil pH 7.24). To assess the impact of water availability on the microbiomes, soil plots were subjected to RF or D conditions, achieved using rainout shelters. Gravimetric moisture content decreased under drought conditions for all soil types, but the reduction was most substantial in Podzol (5.58% RF to 3.85% D; 31-fold% decrease), followed by Luvisol (16.16% RF to 14.34% D; 11.26-fold% decrease) and the lowest in Phaeozem (30.42% RF to 29.46% D; 3.16-fold% decrease).

**Figure 1:**
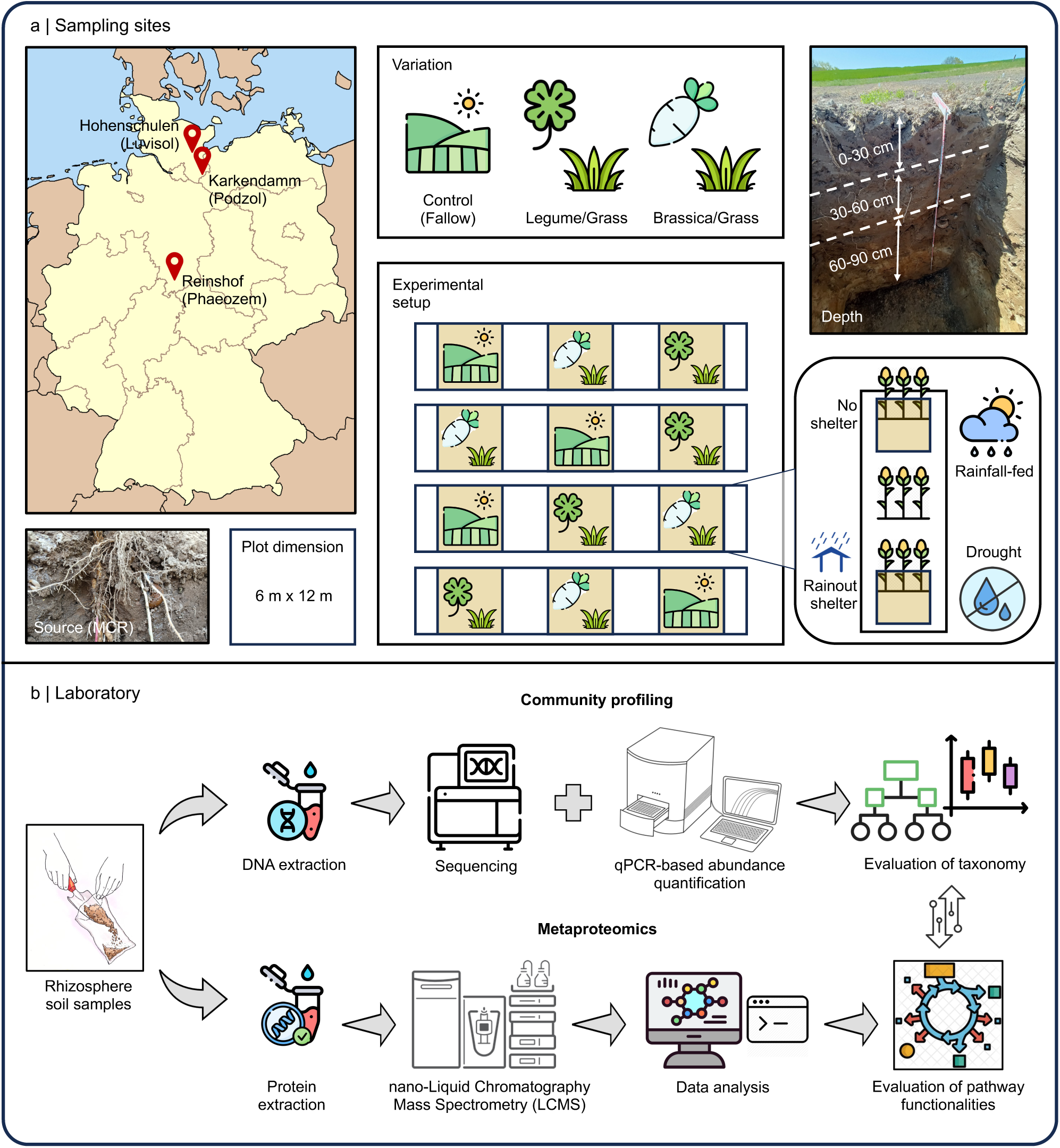
The experimental design for studying reuse of winter cover crop root channels to cultivate maize. **a.** Field sites: samples from Luvisol, Podzol, and Phaeozem in Germany with comparison of bacterial microbiome associated with roots of maize (*Zea mays* L.) in root channels of cover crop [Legume/Grass (*Fabaceae*/*Poaceae*), Brassica/Grass (*Brassicaceae*/*Poaceae*)] or fallow (no cover crops) as control conditions. To replicate drought conditions, plots were covered with rainout shelters. **b.** Community profiling using 16S rRNA gene-based amplicon sequencing, qPCR, and metaproteomics were used to visualise how the composition and functional role of bacterial communities in the maize rhizosphere changed after being exposed to drought.

We estimated the alpha- and beta-diversity of the bacterial communities and detected proteins from the three soil types, sampling depths (topsoil and subsoil) and moisture conditions (D and RF) using 16S rRNA gene amplicon sequencing complemented by qPCR analysis, and metaproteomics. Amplicon sequencing of 150 samples generated 42.8 million high-quality reads, resulting in 117,483 unique amplicon sequence variants (ASVs) assigned to 132 classes within 42 bacterial phyla (Fig. S2). Metaproteomics yielded a total of 2916 proteins belonging to 90 unique functional categories. Bacterial richness, as measured by the Shannon-Wiener index, differed significantly with soil type (Fig. 2a-b, Tables S2a-b,i). Luvisol and Podzol exhibited higher richness (7.080 _Podzol_ and 7.429 _Luvisol_) than Phaeozem (6.956 _Phaeozem_, r = 0.36, \*\*\**p* < 0.001). Bacterial richness was also significantly greater in topsoil compared to subsoil (r = 0.36, \*\*\**p* < 0.001), consistent with previous findings [9]. In contrast, richness did not differ significantly between moisture conditions (7.151_D_ vs 7.159_RF_, r = 0.36, *p* > 0.05), aligning with reports of limited drought impact on soil bacterial alpha-diversity [17, 18]. Proteomic richness also varied significantly among soil types (4.735 _Phaeozem_ vs 4.500 _Podzol_ and 4.839 _Luvisol_, \**p* < 0.05, Fig. 2a-b, Table S2c), but not between D and RF conditions (4.721 _D_ vs 4.661 _RF_, *p* > 0.05, Fig. 2a-b, Table S2h). Bacterial and proteomic evenness did not differ significantly across any of the studied parameters (*p* > 0.05, Fig. 2a, Tables S2d-e,h,j).

**Figure 2:**
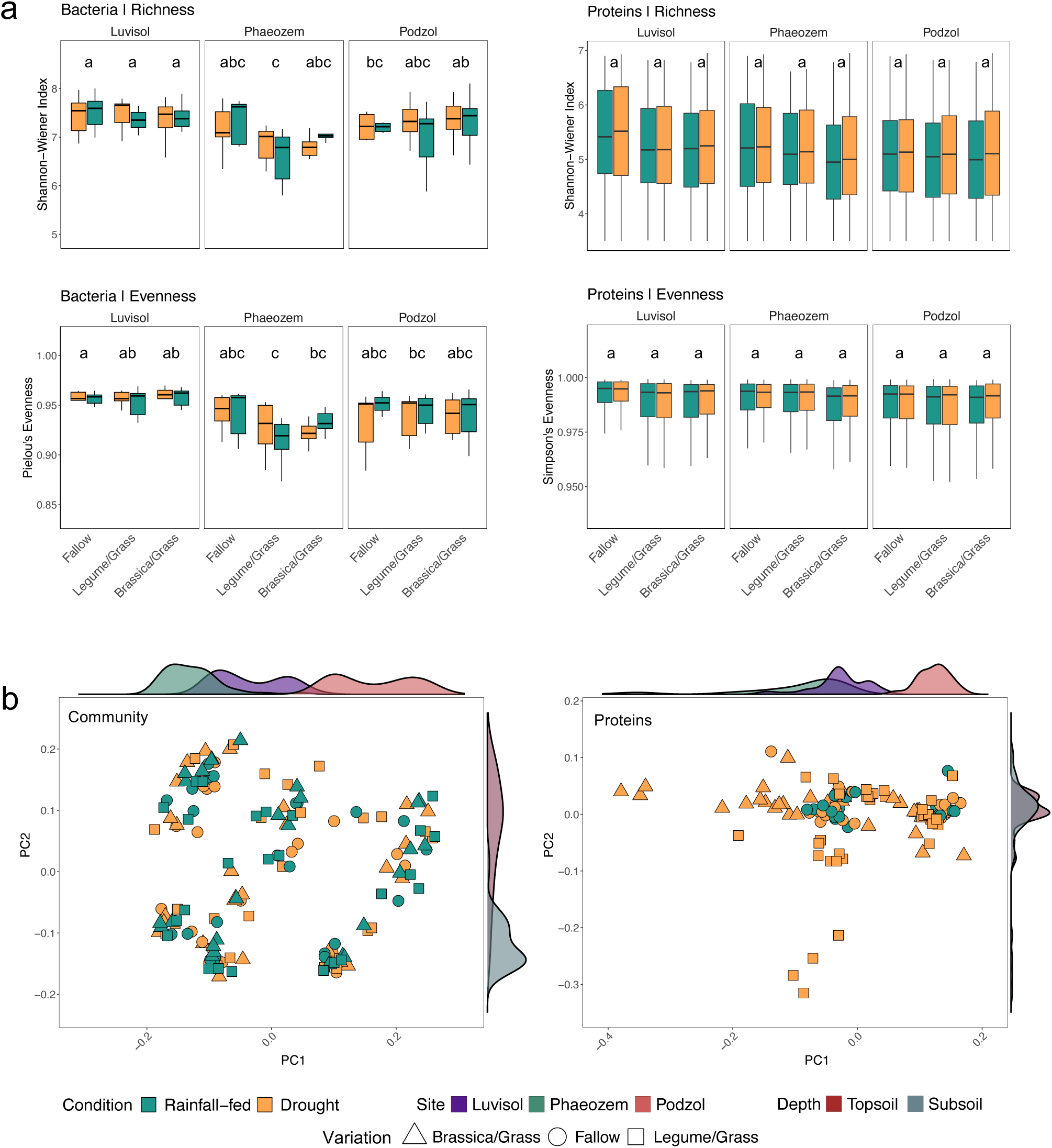
The taxonomic and functional diversity at the different experimental sites under soil moisture conditions. **a.** Alpha diversity richness (Shannon-Wiener index) and evenness (Pielou’s and Simpson’s evenness index) metrices reflecting the distribution of bacterial communities and proteins at the three sampling sites under drought (D) and rainfall-fed (RF) conditions. Pairwise correlation between the variations is shown using a compact letter display representation for the significant differences between the variations, calculated using Tukey’s range test (TukeyHSD); **b.** Bacterial community and proteomic beta-diversity visualised using PCoA ordination based on Jensen-Shannon divergence along the sampling sites and depths for each cover crop variation under soil moisture conditions. Distance metrics were calculated using amplicon sequence variants (ASVs) and unique clusters of orthologous genes (COGs). The different colours of the density plots next to the axes represent the soil types and sampling depths, and peak height and shape abundance. For Luvisol, *n* = 49; for Phaeozem, *n* = 50; and for Podzol, *n* = 51; For soil moisture conditions (drought and rainfall-fed) and depth (topsoil and subsoil), *n* = 150 (statistical details in Supplemental Tables S2f-l)

Enumerations with qPCR revealed a decrease in the average number of 16S rRNA gene copies per gram of soil from 1.6×10^9^ ± 0.5×10^9^ in the topsoil to 3.1×10^8^ ± 2.3×10^8^ in the subsoil (Fig. 3a, Tables 1, S3a). In the topsoil, D conditions in the Podzol were associated with higher copy numbers in re-used root channels from the *Brassicaceae/Poaceae* mixture compared to RF and decreased in the fallow setup under the same conditions. No significant differences in copy numbers were found between D and RF in root channels of the *Fabaceae/Poaceae* mixture, or between topsoil of Luvisol and Phaeozem, regardless of cover crop mixture. In subsoil, cover crop cultivation increased bacterial abundance in the Phaeozem and Podzol compared to fallow, but this effect was not observed in the Luvisol. No differences in bacterial abundance were observed for subsoils between D and RF fallow treatments (Fig. 3a, Tables 1, S2h).

**Figure 3:**
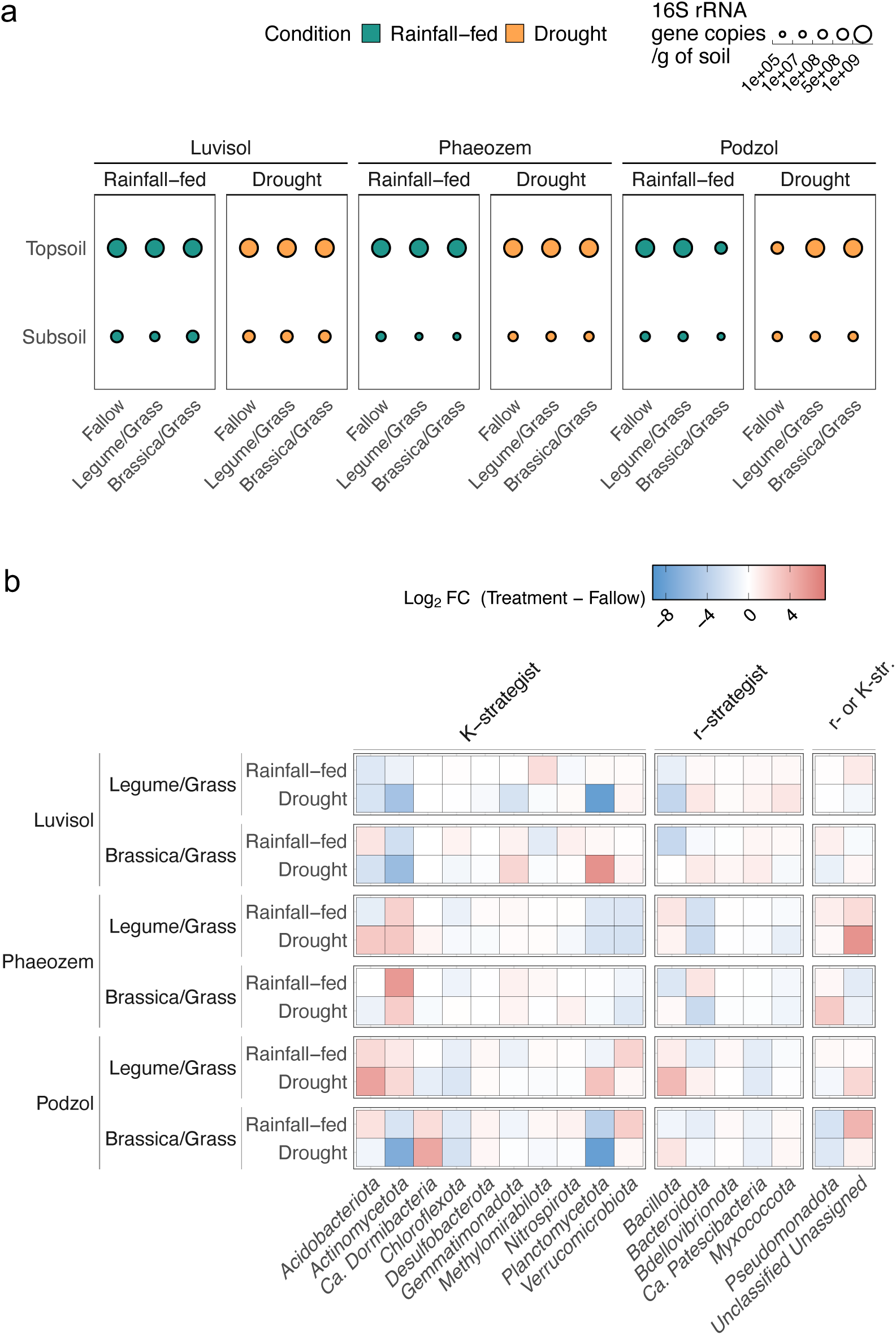
The absolute abundance of bacterial communities in the topsoil and subsoil at three soil types. **a.** 16S rRNA gene copies measured using qPCR as surrogate for absolute bacterial abundances in the soil types (Luvisol, *n* = 49; Phaeozem, *n* = 50; and Podzol, *n* = 51), of each cover crop variation (Fallow, Legume/Grass and Brassica/Grass) and soil moisture conditions (drought (D) and rainfall-fed (RF)). **b.** Relative changes of each bacterial phylum (categorised as *K*-strategist, r-strategist and r or *K*-strategist) in reused cover crop root channels under D and RF conditions. Only phylotypes displaying significant increase or decrease in abundance are shown.

**Table 1:**
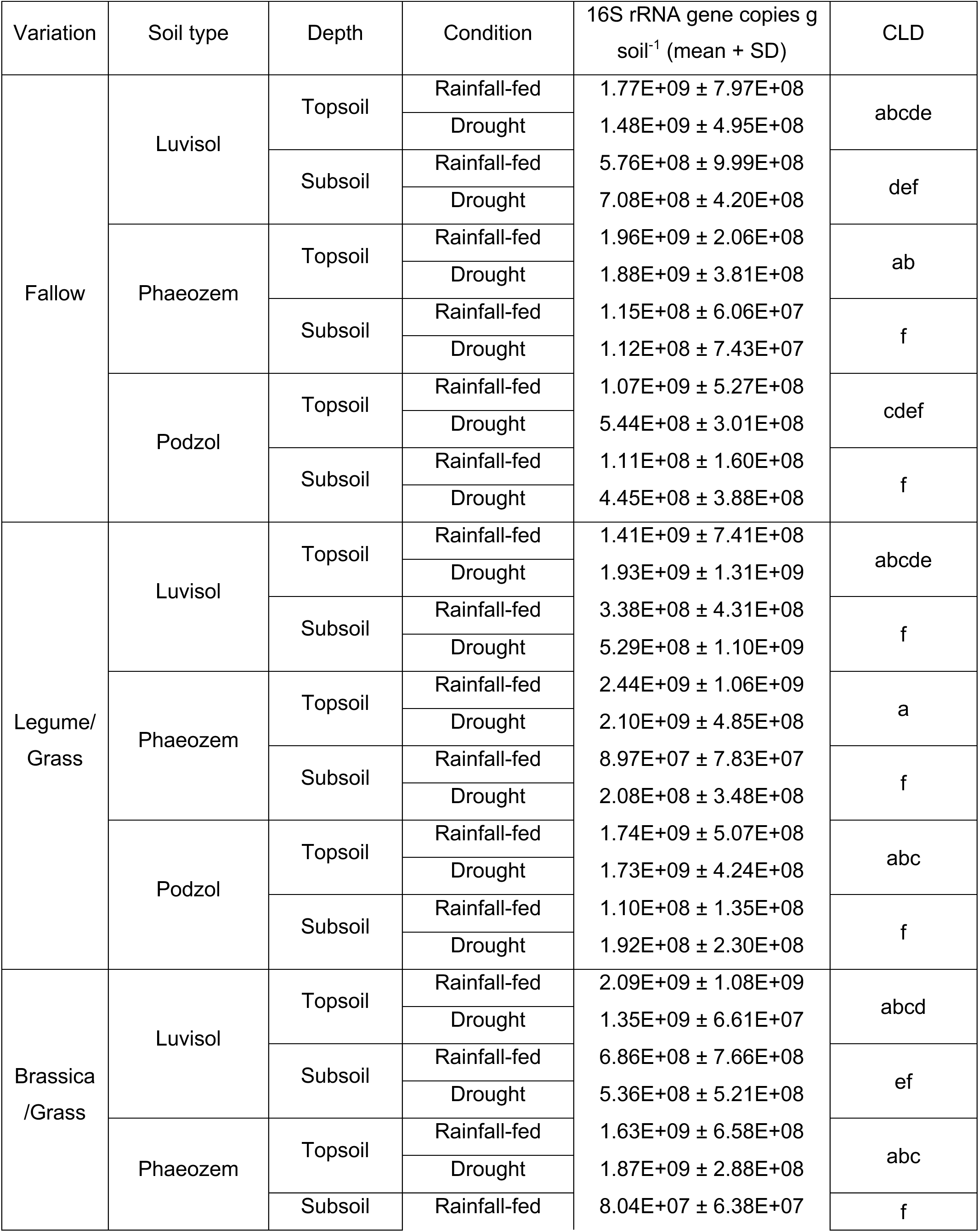
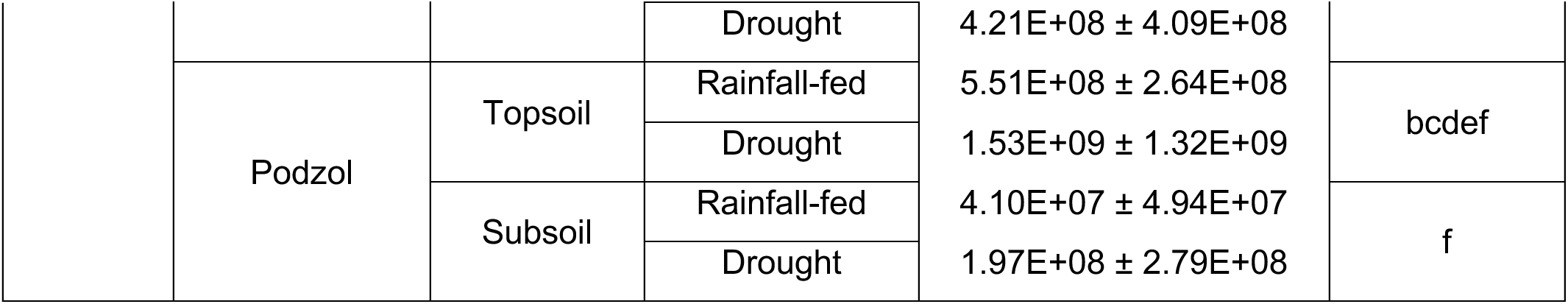
Summary of 16S rRNA gene copies per gram soil for different parameters in our study. We quantified the mean and SD values of 16S rRNA gene copies per gram soil in soils from the reused cover crop root channels for all study parameters. The significance of each category is represented using compact letter display (CLD) calculated after conducting MANOVA on the gene copies under categories of soil type, cover crop variations and soil sampling depths. The values are provided in the Supplemental Tables S2f-h, S3a.

While alpha-diversity did not differ between D and RF conditions, beta-diversity varied significantly with soil types and sampling depths for both 16S rRNA gene amplicon and metaproteome data, as calculated using Jensen-Shannon divergence coupled with PERMANOVA and Tukey’s post-hoc tests (Fig. 2b, Table S2a). We describe the shifts in beta-diversity using abundances of selected phyla (Tables S2f-g, S4). The changes were particularly pronounced for *K*-strategists (Fig. 3b, 4a, Table S3b). In cover crop root channels reused by maize roots, *Acidobacteriota* significantly increased under D in the acidic Podzol but decreased under D in the Luvisol (Log2FC: 2.13_Podzol_ vs -0.20_Phaeozem_ vs -2.33_Luvisol_). *Actinomycetota*, another *K*-strategist, increased in the neutral-to-basic Phaeozem under D (-1.15_Podzol_ vs 3.47_Phaeozem_ vs 0.09_Luvisol_). *Planctomycetota* decreased significantly in Phaeozem but increased in Luvisol and Podzol (3.47_Podzol_ vs -1.33_Phaeozem_ vs 5.45_Luvisol_) under D, whereas *Pseudomonadota* increased in all three soil types under D (0.13_Podzol_ vs 1.18_Phaeozem_ vs 0.44_Luvisol_), which is similar to previous findings under dry conditions [19]. In contrast, *Chloroflexota* and *Verrucomicrobiota* decreased under D (-0.14*_Chloroflexota_* and -0.63*_Verrucomicrobiota_*) compared to RF conditions. Changes in the abundance of 11 out of the 132 identified bacterial classes differed from that of their respective phylum (e.g. *Symbiobacteria* of *Bacillota* and UBA8108 of *Planctomycetota*) during drought, exemplifying the notion that members of a bacterial phylum do not always share similar ecophysiological traits [20]. *Candidatus* Dormibacteria, a newly-characterised taxon that is also considered a class of *Chloroflexota* and known to favour dry, nutrient-poor soils [21], was abundant in sandy Podzol and in the deeper subsoil (10-fold increase in abundance in Podzol compared to Luvisol and Phaeozem; 7.3-fold increase in subsoil as compared to topsoil, Fig. 4b, Table S3b).

**Figure 4:**
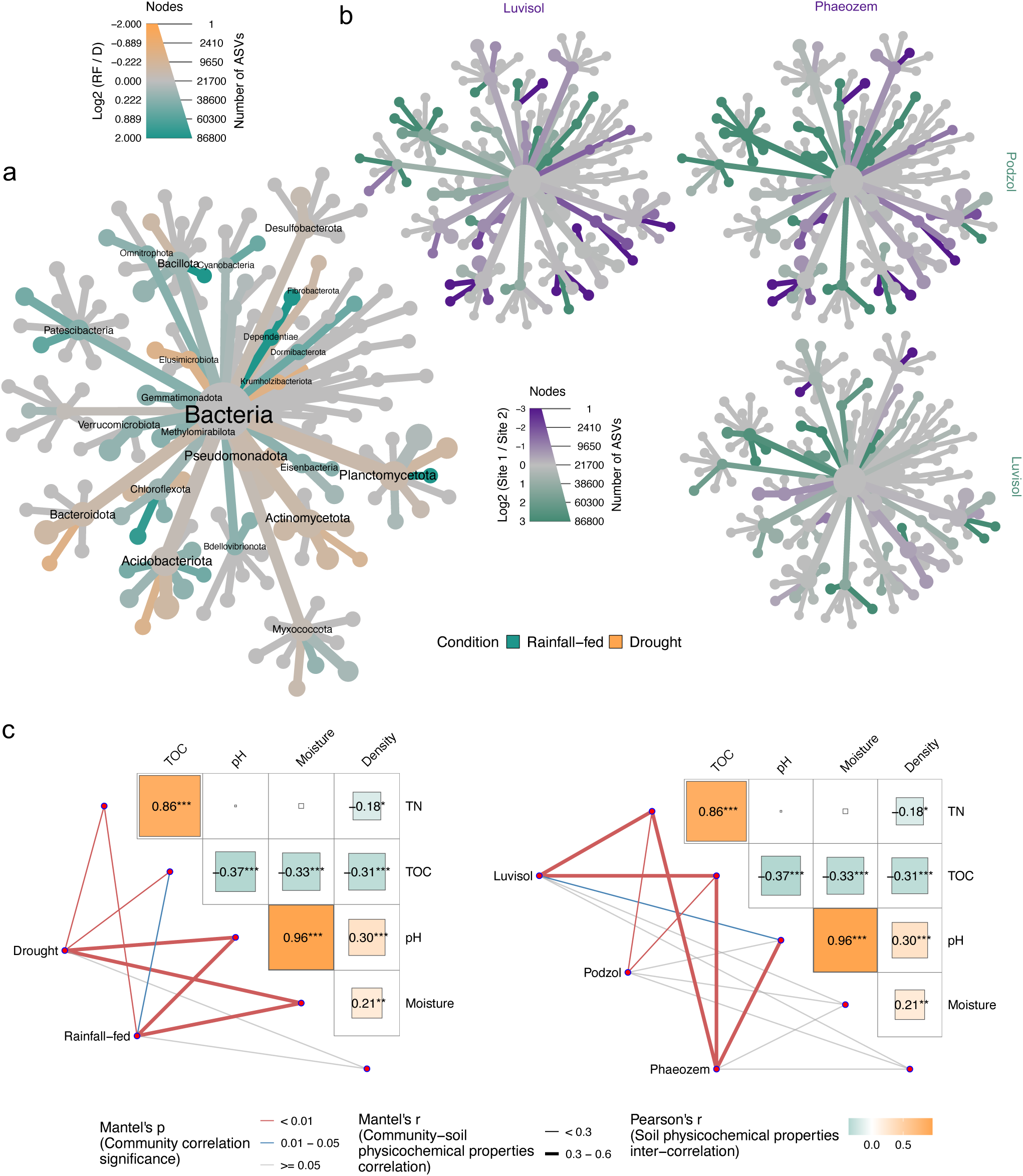
Correlations among soil physicochemical properties and bacterial community framework for soil types and moisture conditions. Taxonomic heat trees represent the differential abundance of the bacterial phylotypes between two parameters of (**a**) soil moisture content [drought (D) vs rainfall-fed (RF)], and (**b**) soil types (Luvisol, Phaeozem and Podzol). The yellow-ochre colour indicates increases under D conditions and green colour under RF conditions, **c**. Partial Mantel’s test: the matrix represents the correlation of the soil physicochemical parameters, the lines the correlation between the community distance matrix, and the thickness the strength of the correlation and the colour indicates the significance; \**p* < 0.05, \*\**p* < 0.01, \*\*\**p* < 0.001.

### Comparing soil physicochemical properties

To investigate the influence of soil physicochemical properties on the bacterial microbiome in the maize root channels, we correlated the parameters pH, soil moisture content, TOC, TN, and bulk density with the 16S rRNA gene amplicon data using Partial Mantel’s test [22] (Fig. 4c) and confirmatory factor analysis (CFA) [23] (Fig. S3). Partial Mantel tests revealed significant relationships between community structure and pH, soil moisture content, C and N (\*\**p* < 0.01, Fig. 4c, 5, Table S5a). The strength of the correlation between community composition, moisture, and pH decreased under D as compared to RF (r_pH, D_ = 0.473 vs r_pH, D_ = 0.525, r_H_2_O, D_ = 0.416 vs r_H_2_O, RF_ = 0.483, Table S5b).

CFA modelling further supported that pH (0.71_CFA_), soil moisture content (0.07_CFA_), and bulk density (0.10_CFA_)) were key physicochemical factors influencing community composition, with a strong connectedness between the three factors (Fig. S3, \*\*\**p* < 0.001, Table S5c, S6). The modelling also indicated that cover crop roots provide increased access to total soil organic carbon (Fig. S4-5, Table S5c) [24, 25]. The effect of blocking precipitation via rainout shelters on the bacterial communities diminished with depth (r_Topsoil_ = 0.704, r_Subsoil_ = 0.581, Fig. S6-7, Table S5d), potentially due to reduced drought stress and more stable water content in the subsoil. At the community level, *Cyanobacteria* and *Planctomycetota* abundances positively correlated with TOC and TN (Fig. 5, Table S5e), consistent with previous reports [26, 27]. Conversely, *Chloroflexota*, *Methylomirabilota* and *Nitrospirota* exhibited negative correlations with TOC and TN across all three soil types (Fig. 5, Table S5f).

**Figure 5:**
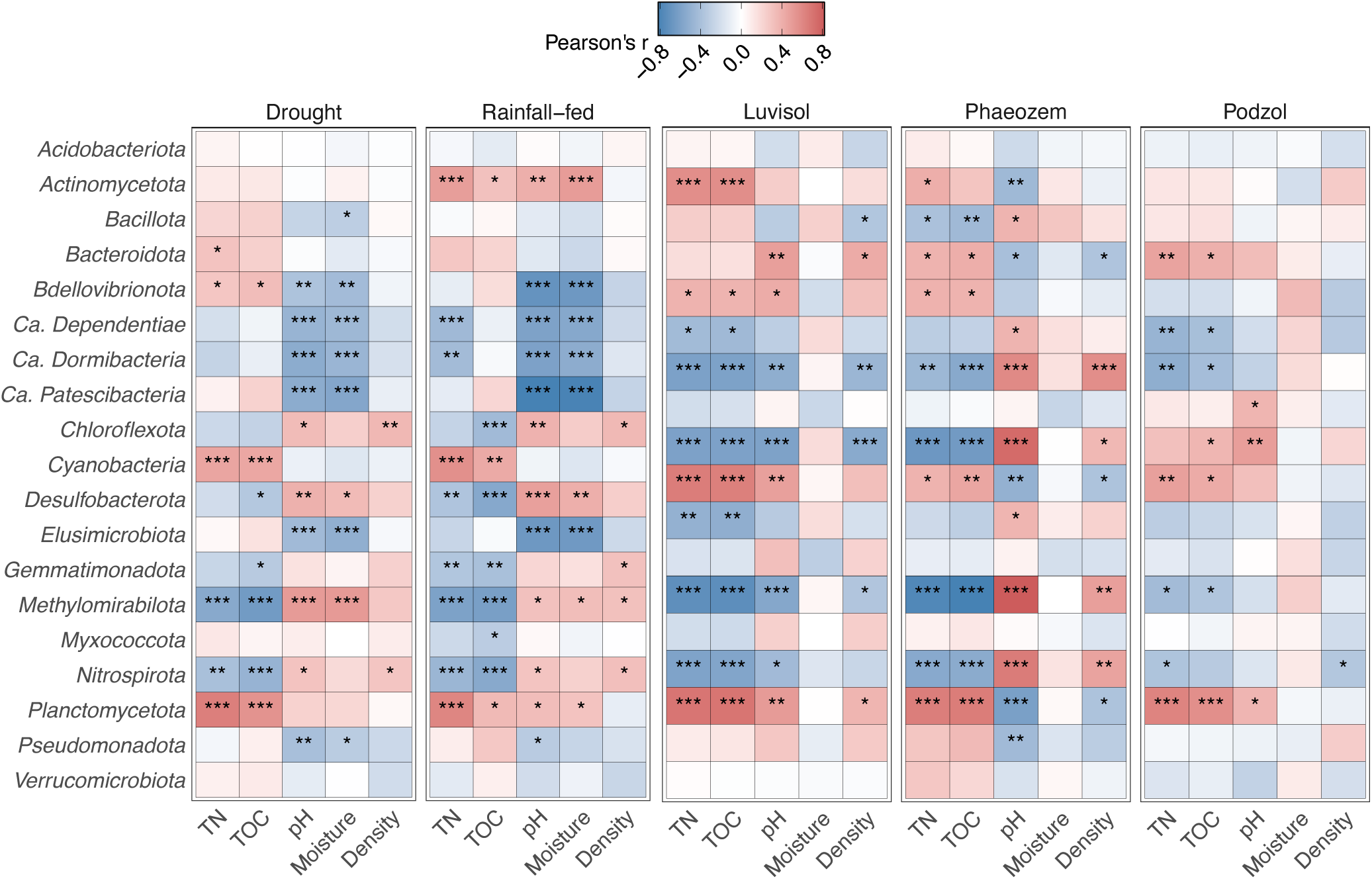
Correlation of individual bacterial communities and the soil physicochemical properties. The correlations between each bacterial phylum with the physicochemical properties (TN, TOC, pH, Moisture and Density) have been estimated using Partial Mantel’s test. Pearson’s correlation values and the significance estimation for each physicochemical protein and each phylum are provided in the Supplemental Tables S5e-f. The significant correlations are represented by asterisks inside the heatmap tiles; \**p* < 0.05, \*\**p* < 0.01, \*\*\**p* < 0.001.

Lower TOC and TN in the subsoil appeared to favour *Chloroflexota*, *Methylomirabilota* and *Nitrospirota*, potentially allowing them to outcompete other phyla (Fig. 5, S2, Tables S1, S5e-f, S6). Similarly, members of rare taxa such as *Ca.* Dependentiae and *Ca*. Patescibacteria (also known as Candidate Phyla Radiation) increased relatively in the subsoil compared to the topsoil in reused cover crop root channels (Fig. 5, S2, Tables S1, S5e-f, S6). Additionally, the correlation between *Methylomirabilota* and *Ca*. Patescibacteria with moisture was stronger under D than under RF conditions (Fig. 5).

### Functional response of root channel-residing bacteria towards drought

Metaproteomic analysis revealed the functional response of bacterial communities within maize root channels to D conditions. Identified proteins were assigned to taxonomic groups with custom generated databases from UniProt containing all the reference proteomes for those communities identified in 16S rRNA amplicon sequencing. The corresponding pathways were categorised using the Kyoto Encyclopaedia of Genes and Genomes (KEGG) [28]. Protein group abundances varied significantly with soil type, depth, and cover crop variation, and were consistent with 16S rRNA gene amplicon sequencing results (Table S7a). Samples from the Podzol consistently exhibited the highest abundance, followed by Phaeozem and Luvisol (Table S7b). More protein groups were found in topsoil compared to subsoil across all soil types (582_Podzol-Topsoil_ > 455_Phaeozem-Topsoil_ > 259_Luvisol-Topsoil_; 330_Podzol-Subsoil_ > 262_Phaeozem-Subsoil_ > 188_Luvisol-Subsoil_) and in D compared to RF samples (443_D, avg._ > 355_RF, avg._). Protein expressions in D vs RF increased in the Podzol and the Phaeozem in reused cover crop root channels (0.677_Podzol-Log2FC_, 0.576_Phaeozem-Log2FC_), but decreased in the Luvisol (-0.665_Luvisol-Log2FC_), and did not significantly change in fallow, assessed by Wilcoxon and multivariate analysis of variance (MANOVA) tests (Tables S7c-d). Root channels from the *Brassicaceae*/*Poaceae* mixture exhibited the greatest variability compared to *Fabaceae*/*Poaceae* and fallow, possibly reflecting differences in soil physicochemical properties of their pore walls such as water content, bulk density, and C and N availability. Detailed metadata of all identified proteins and the associated biochemical pathways are provided in Table S8.

Volcano plot analysis identified differentially expressed enzymes under D stress, with a greater number of upregulated proteins in the root channels of *Brassicaceae/Poaceae*, especially in the drought-affected Luvisol and Podzol soils (Fig. 6, Table S9). The upregulated enzymes were primarily involved in glucose and pyruvate metabolism, the glyoxylate shunt of the citric acid cycle (TCA), amino acid synthesis, the methionine cycle, and the transsulfuration pathway (Fig. 7a, Tables S9, S10a). Specifically, several glycolytic enzymes, including aldolase, enolase, phosphoglucose isomerase and triose phosphate isomerase (TIM), were significantly upregulated under drought (\**p* < 0.05, Tables S9, S10b). Upregulation of phosphoenolpyruvate carboxykinase (PEPCK) (\*\*\**p* < 0.001, Tables S9, S10a) likely facilitated the use of oxaloacetate for pyruvate synthesis and subsequent gluconeogenesis. Expression of enzymes from the pentose phosphate pathway, 6-phosphogluconate dehydrogenase (6PGDH) and glucose-6-phoshate dehydrogenase (G6PDH), increased under drought (\*\*\**p* < 0.0002), matching previous studies with rice, tomato and soybean roots but without cover cropping [29-31]. Elevated expression of pyruvate dehydrogenase complex (PDC) proteins and acetyl-CoA synthetase under D conditions suggest increased synthesis of acetyl-CoA from pyruvate, acetate, and fatty acid oxidation, potentially promoting the glyoxylate shunt, a TCA cycle variant favoured under such conditions [32] (Fig. 7a). Upregulation of the glyoxylate shunt was evidenced by significantly increased abundance of malate synthase, which converts glyoxylate to malate. Catalase-peroxidase (CAT-PER) and superoxide dismutase (SOD) abundances were higher in the Luvisol and the Podzol compared to the Phaeozem, suggesting enhanced reactive oxygen species (ROS) scavenging (\*\*\**p* < 0.001, Fig. 7b). In the Podzol, the majority of enzymes involved in the methionine cycle and transsulfuration pathway were also overexpressed (\*\**p* < 0.01). These two pathways produce glutathione, needed for facilitating CAT-PER based breakdown of ROS (Fig. 7b, Tables S9, S10a). Enzymes contributing to amino acid synthesis, including 1-pyrolline-5-carboxylate dehydrogenase, aldehyde dehydrogenase, branched-chain transaminase, and glutamate-gamma semialdehyde dehydrogenase, were also more abundant under D conditions across all three soil types (Fig. S8). In the N cycle, glutamine synthetase was the only enzyme that showed a differential expression, specifically an upregulation in the Podzol and downregulation in the Luvisol and Phaeozem under D compared to RF conditions with *Brassicaceae/Poaceae* as the winter cover crops. (Table S9).

**Figure 6:**
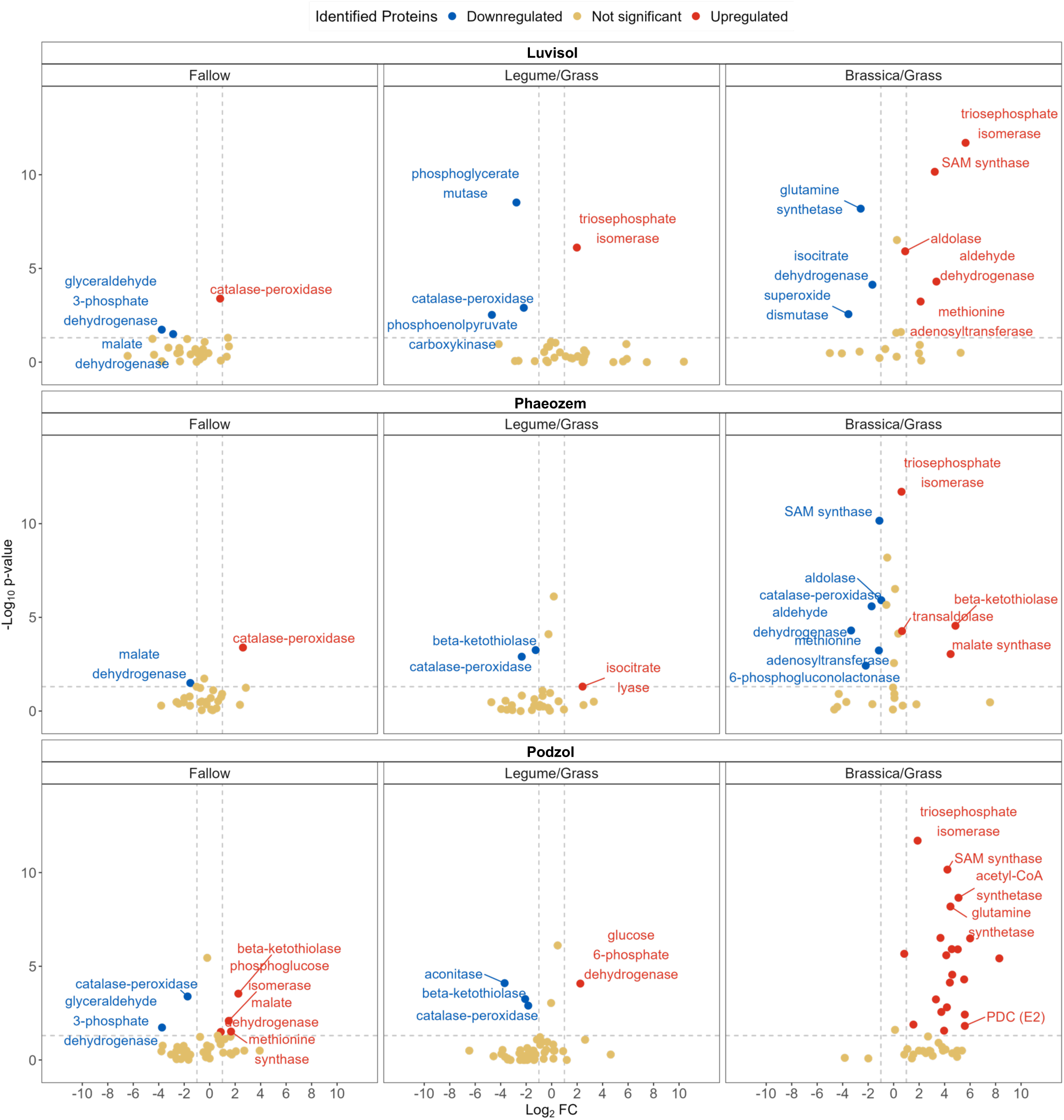
Upregulated and downregulated proteins under drought. A volcano plot representation for the upregulation and downregulation of the identified proteins under drought conditions at the three sampling sites (Luvisol, Phaeozem and Podzol) for the cover crop variations (Fallow, Legume/Grass and Brassica/Grass). The *x*-axis represents the Log2 fold-change and the *y*-axis represents the -Log_10_ p-value according to Mann-Whitney U test. The proteins having Log2 fold-change (FC) values < 0.6 or > -0.6 and p > 0.05 were classified as ‘Not significant’. Log2FC > 0.6 and p < 0.05 were the ‘Upregulated’ proteins and Log2FC < 0.6 and p < 0.05 were the proteins ‘Downregulated’. Names of several identified proteins are not shown to avoid crowdedness. The information for all proteins is provided in the Supplemental Table S9.

**Figure 7:**
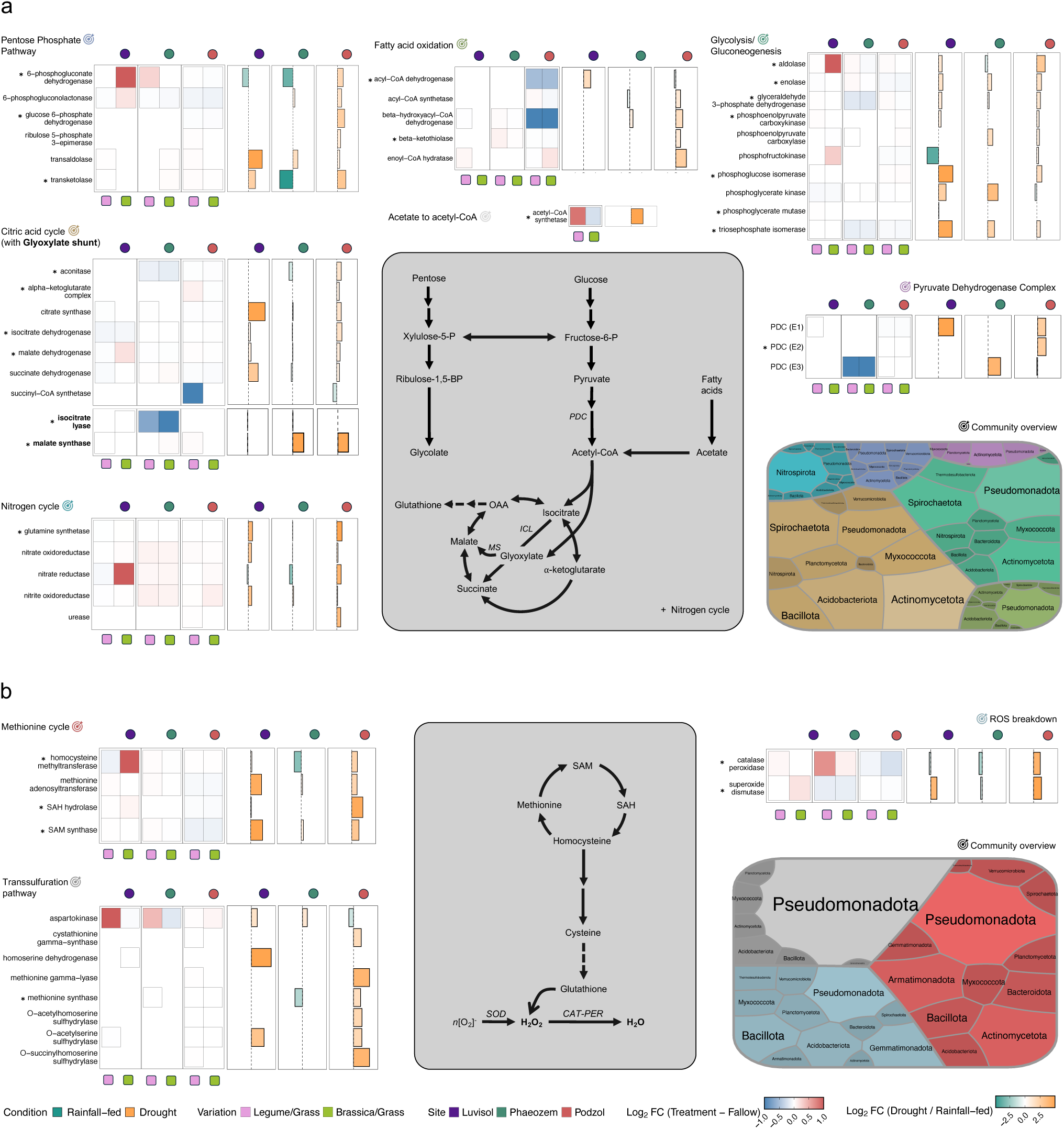
Overview of the functional pathways affected by drought in three soils after cover crop root-channel-reuse. Pathways with contributions to (**a)** catabolism to (**b)** anabolism. Heatmaps are showing the net expression change based on measured protein LFQ intensities for each protein identified. Positive logarithmic values of the difference are denoted by red (higher in re-used root channel than in bulk soil rhizosphere) and negative values by blue (lower in re-used root channel than in bulk soil rhizosphere). The bar charts represent whether the proteins were differentially expressed under drought. The small circles and boxes at the top and below the heatmaps and bar charts code for the soil types and cover crop mixtures respectively. Tree maps represent the bacterial phyla identified as pathway source. The significance of the changes in protein expression under drought conditions at each site was calculated using a four-way ANOVA model, significant changes in expression are marked with an asterisk (\**p* < 0.05) (Supplemental Table S10a consists of a list of proteins which represented significant changes across the parameters of study).

Comparing relative protein expression between cover crop mixtures and fallow treatment revealed greater variability and upregulation in topsoil under D versus RF conditions, while subsoil expressions were similar under both conditions (Fig, S9, Tables S7e, S9). Protein expression was significantly higher in topsoil compared to subsoil (2.11-fold increase, Table S7e). The identified bacterial proteins were predominantly from *K*-strategist. In topsoil, the phyla *Acidobacteriota*, *Actinomycetota*, *Armatimonadota*, *Myxococcota*, *Planctomycetota*, *Pseudomonadota*, and *Verrucomicrobiota* contributed the highest abundances of identified proteins. In contrast, *Bacillota*, *Chloroflexota*, *Methylomirabilota*, and *Thermodesulfobacteriota* were the dominant sources of identified proteins in the subsoil. Enzymes of the glyoxylate shunt, cysteine and methionine metabolism, and ROS breakdown were mostly from *Methylomirabilota*, *Pseudomonadota*, *Spirochaetota* and *Verrucomicrobiota*.

The expression of proteins indicative of general metabolic activity, including ribosomal proteins, chaperonins, transmembrane proteins, and peroxidases, were significantly higher in reused cover crop root channels compared to fallow (\*\*\**p* < 0.001) (Fig. 8, Tables S11a-b). Under D conditions, the abundance of these proteins increased by multiple orders in root channels as compared to RF (21.9-fold increase in chaperonins; 16.9-fold increase in heat shock proteins and 17.9-fold increase in ribosomal proteins), with ATP synthase being the most abundant protein detected, which increased by 0.2-fold under D against RF. The majority of these proteins was sourced from the rare phyla *Armatimonadota* and *Ca.* Latescibacterota.

**Figure 8:**
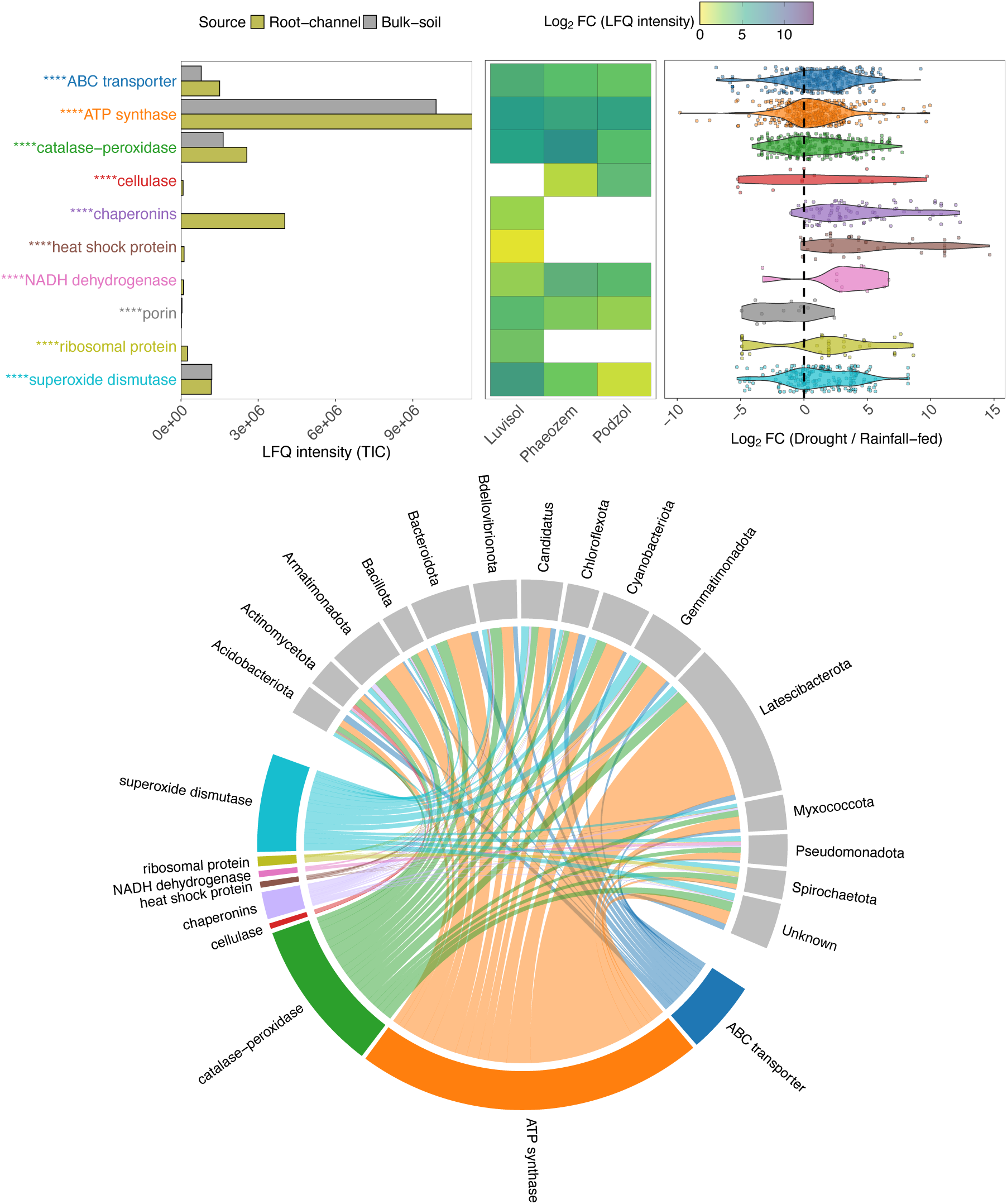
Estimation of bacterial activity. Chord diagram showing abundance changes of predominant proteins (chaperonins, ribosomal proteins, ROS-regulators, membrane proteins such as porins, ATP synthase). Substantial abundance changes indicate high translation activity, and hence overall metabolic activity. Protein presence in the three different soil types is represented by the heatmap, followed by the comparative abundance under drought conditions as compared to rainfall-fed conditions, represented as violin plot. proteins showing significant changes in expression are marked with an asterisk (\**p* < 0.05).

## Discussion

This study investigated the impact of drought on bacterial communities and expressed biochemical pathways in winter cover crop root channels reused by maize. The reuse of cover crop root channels facilitates maize root propagation into the subsoil, likely by reducing penetration resistance and potentially by increasing nutrient availability in the channel wall accessible to the bacterial community beneficial to maize [9]. Here, increased nutrient availability was evidenced by greater copy numbers of 16S rRNA gene in reused cover crop root channels, particularly from *Brassicaceae/Poaceae* in Podzol, compared to maize root channels in fallow. Building upon previous research examining drought effects on soil biogeochemistry and microbial communities [33-35], our findings reveal relationships between type of cover cropping, soil type, moisture availability, and bacterial microbiome dynamics and function.

Bacterial alpha-diversity and protein distributions differed significantly among soil types (Luvisol > Podzol > Phaeozem), but were not affected by soil moisture conditions. The observed higher richness in drought-impacted Luvisol and Podzol soils may reflect the legacy of decade-long water scarcity in the region [36], consistent with recent reports of increased bacterial richness under prolonged drought [37]. We observed a greater prevalence of *K*-strategists (e.g., *Acidobacteriota*, *Actinomycetota*, *Planctomycetota*) across our samples, consistent with the resilience of slow-growing bacteria to environmental stress [38-40]. These bacteria are better adapted to resource-limited conditions and can maintain activity under adverse conditions. Conversely, the decline in oligotrophic phyla (*Chloroflexota*, *Ca.* Dependentiae, *Methylomirabilota*, *Ca.* Patescibacteria) in the topsoil may be linked to increased O_2_ levels in root channels under D conditions (Fig. 3b, 4a, S6, Tables S3c-d). These phyla generally thrive in oxygen-deficient environments such as water-filled pore space [9, 41-44]. The increased abundance of *Ca.* Dependentiae and *Ca.* Patescibacteria in the subsoil following cover crop cultivation, potentially due to exploiting stimulated rhizodepositions [45] (Table S3d). This is likely facilitated by the ability of these bacteria to prosper under oxygen-rich conditions, aligning with reported inverse relationships between their abundance and with soil moisture [46]. The acidic-to-slightly acidic conditions of the Podzol and the Luvisol favoured the proliferation of *Acidobacteriota*, *Actinomycetota*, *Armatimonadota*, and *Bacteroidota*. The latter two phyla exhibit tolerance to lower soil moisture, potentially contributing to their increased abundance under D conditions [47, 48]. Furthermore, we observed a drought-induced shift towards phyla adapted to nutrient-poor conditions, including *Ca.* Dormibacteria, *Methylomirabilota*, and *Nitrospirota*, at the expense of *Chloroflexota*. While the drier channels during drought may have greater availability of O_2_, reduced water content limited nutrient diffusion, favouring phyla better adapted to nutrient scarcity. Phaeozems, characterized by high soil organic carbon content, supported the proliferation of organic matter decomposers such as *Actinomycetota* [49], which thrive across a broad pH range [50]. Conversely, conditions in Phaeozems may lead to *Planctomycetota*, which are adapted to nutrient-deficient environments [51], being outcompeted by other bacteria.

Physicochemical properties in the rhizosphere of the re-used root channels, including pH, moisture content, total nitrogen, and organic carbon are key determinants of soil quality and bacterial community structure. While previous studies have demonstrated variable relationships between these properties, we observed a positive correlation between pH and soil moisture, potentially linked to changes in C metabolism and proton release during nitrification as soils dry [52-54]. The mobilisation and transport of organic matter from one part of the soil profile to another is only known for the Podzol due to the acidic nature of the soil. The drying up of water from the soil pores hinders organic matter mobility and increases their accumulation, which may explain the negative association between pH and organic carbon [55, 56]. Drought-induced increases in soil aeration also exposes previously inaccessible carbon sources to aerobic bacteria, probably initiating its mineralization [34, 57].

To investigate these relationships further, we measured the abundance of proteins across root channels from the Luvisol, Podzol, and Phaeozem soils under D and RF conditions using metaproteomics. Our analysis revealed distinct responses to drought, with upregulation of glycolytic and pentose phosphate pathway enzymes observed in the drought-affected Luvisol and Podzol, but not in the Phaeozem, which maintained higher moisture levels with higher TOC. The observed increased expression of glycolytic enzymes in the Luvisol and the Podzol likely reflects increased substrate availability due to enhanced aeration in the drier soil leading to the utilisation of previously sequestered TOC [15, 58]. The increased bacterial activity was further manifested by elevated levels of the pyruvate dehydrogenase complex, acetyl-CoA synthetase, and enzymes for β-oxidation of fatty acids. Consistent with these findings, ^13^C labelling studies have reported an accumulation of two-carbon compounds, such as acetate, in drought-affected soils [59, 60]. Increased availability of acetate and metabolism of fatty acids would also explain the higher abundance of enzymes of the glyoxylate cycle, malate synthase and isocitrate lyase [61]. Furthermore, there was an increased abundance of carbohydrate-active enzymes (CAZymes), particularly β-glucosidase, from *Bacteroidota* in *Brassicaceae/Poaceae* root channels of the Luvisol (Fig. S10).

The pentose phosphate pathway also contributes to NADPH generation needed for maintaining cellular redox homeostasis mechanisms, specifically in the glucose-6-phosphate dehydrogenase (G6PDH) and 6-phosphogluconate dehydrogenase (6PGDH) reactions [62]. Both enzymes were upregulated under D conditions. The NADPH is utilized by enzymes such as catalase (CAT), peroxidase (PER), glutathione peroxidase (GPX), and superoxide dismutase (SOD) to mitigate ROS-induced stress, which likely occurs during drought within the well-aerated root channels. Furthermore, increased abundance of aldehyde dehydrogenase, branched-chain transaminase, and glutamate-gamma-semialdehyde dehydrogenase suggests enhanced amino acid synthesis, particularly glutamate and proline, which function as osmoprotectants for bacteria living in shrinking water films during drought. The expression of these pathways was particularly pronounced in the topsoil, likely due to its direct exposure to drought stress, while the subsoil exhibited greater stability in protein expression. The observed upregulation of enzymes involved in the glyoxylate cycle and ROS scavenging suggests that aerobic bacteria, particularly members of the *Pseudomonadota*, *Spirochaetota*, and *Verrucomicrobiota* phyla, respond to these stresses stronger than other phyla. These enzymes can, therefore, represent plausible indicators of drought stress in soil profiles.

The increased abundance of proteins associated with membrane function, ribosomes, and electron transport following maize root colonization of former cover crop channels suggests heightened bacterial activity. This observation supports the previous finding that root channels created by cover crop roots facilitate the formation of microbial hotspots within the maize rhizosphere [10]. Notably, the significant contribution of rare taxa – bacterial phylotypes with a relative abundance below 0.01%, including members of *Armatimonadota* and Ca. Latescibacterota, indicates increased and diversified microbial activity. This diversification likely stems from the complex composition of organic matter within these channels, a mixture of decaying root detritus and microbial necromass from the previous cover crop rhizosphere, combined with fresh root exudates from maize. This complex substrate base likely enhances access to subsoil resources [9]. The cover crops mixture with *Brassicaceae* and *Poaceae* exhibited the most pronounced upregulation of these pathways, particularly in rapidly draining soil types like the Luvisol and the Podzol. *Brassicaceae* are known for establishing deep root channels and efficiently exploiting subsoil resources, even during short-term winter cover cropping. Consequently, the microbiome inhabiting these channels may be pre-adapted to utilize subsoil resources, potentially buffering maize against drought-induced stresses.

Understanding the full complexity of microbial interactions within root channels remains a significant challenge. The walls of re-used cover crop root channels represent an open system characterized by diverse microhabitats and influenced by fluctuating physicochemical factors. We observed that certain bacterial phylotypes, such as *Symbiobacteria* (*Bacillota*) and UBA8108 (*Planctomycetota*), exhibited abundances inconsistent with broader patterns of their respective phyla, illustrating the nuanced dynamics and potentially different ecophysiological traits of these microorganisms. Integrating community profiling with metaproteomics, coupled with the continued expansion of reference databases, will be crucial for resolving these complexities. Overall, our approach successfully generated a substantial proteomic dataset, providing a functional overview of biochemical pathway shifts under D conditions and revealing the key roles played by specific bacterial community members. Our findings demonstrate that drought stress elicits a coordinated bacterial response in the rhizosphere of maize reusing cover crop root channels, characterized by upregulation of the glyoxylate cycle and enzymes involved in ROS scavenging. The observed responses are particularly pronounced in the topsoil and modulated following *Brassicaceae* and *Poaceae* cover crops. These results highlight the potential for leveraging cover cropping strategies to mitigate the impacts of drought on agricultural systems.

## Materials and Methods

### Crop cultivation and sampling regimes

Crops were grown in three agricultural fields – at experimental estates Hohenschulen of the Christian-Albrechts-University of Kiel (Achterwehr, Germany, 54°18’44” N, 9°59’46” E), Karkendamm of the Christian-Albrechts-University of Kiel (Bad Bramstedt, Germany, 53°55’52” N, 9°55’15” E) and Reinshof of the Georg-August-University of Göttingen (Rosdorf, Germany, 51°29’05” N, 9°53’34” E). The cash crop in this study was maize (*Zea mays* L.). Plots without cover crops during the winter (bare fallow) were established as control and compared against two cover crop mixtures on distinct plots. The cover crops were shallow- and deep-rooting *Brassicaceae* (*Brassica napus* L., rapeseed, shallow-rooting; *Raphanus sativus* L. var. *oleiformis*, oilseed radish, deep-rooting); *Fabaceae* (*Trifolium repens* L., white clover, shallow-rooting; *Trifolium pratense* L., red clover, deep-rooting); and *Poaceae* (*Lolium perenne*, perennial ryegrass, shallow-rooting; *Festuca arundinaceae*, tall fescue, deep-rooting). The mixtures were grown as a combination of shallow- and deep-rooting cover crops of *Brassicaceae*, *Fabaceae* and *Poaceae,* complementing the niche complementary principle, which has been reported to allow polycultures to over-yield when plants compete for resources [63]. All the cover crops were sown in October 2022 and grew until May 2023 in distinct randomised plots with four replicates of each variation. In May 2023, herbicides (Roundup, Bayer AG, Leverkusen, Germany) were used to kill all cover crops, and subsequently, maize was sown in the same plots with the cover crop variations in addition to the fallow profiles. Maize was grown in the field from May to August 2023 (Fig. 1). To compare drought-like conditions against normal conditions, we artificially induced dry conditions using interrow rainout shelters. These specific type of rainout shelters covered the area between the maize rows, largely removing the throughfall and restricting water infiltration to stemflow. The profiles were covered with those shelters after the maize exceeded 30 cm height and maintained until sampling was done around the flowering and reproduction stage of maize. The field design was structured as a randomized block design with the four blocks representing the four replicates and the ambient (natural rainfall) and drought (with interrow rainout shelters) plots being always paired in direct proximity to each other.

Prior to each sampling, the soil profile was excavated 40 cm forward to obtain a fresh profile and fresh maize root system and to prevent any contamination by the neighbouring soil. To compare the difference inflicted by drought on microbial communities in the reused root biopores of cover crops, we collected soil samples from a vertical soil profile from the topsoil (0-30 cm) and the subsoil (30-60 and 60-90 cm) for three variations: 1) fallow, where maize was grown on profiles without any pre-cultivated winter cover crops, 2) *Brassicaceae*/*Poaceae*, where maize grown in the plots with *Brassicaceae* and *Poaceae* grown during winter, 3) *Fabaceae*/*Poaceae*, where maize grown in the plots with *Fabaceae* and *Poaceae* grown during winter. The collected samples were categorised under two specific conditions: 1) maize roots growing in the cover crop root biopores (MCR) from those profiles subjected to drought (D), and 2) maize roots growing in the MCR in the profiles under rainfall-fed conditions (RF). Areas on the profile where fresh white maize root and dark brown cover crop root residues completely overlap were seen as MCR regions. MCR samples were soils taken within 2 mm of the root overlap region. Since fallow was defined as maize roots grown in the profiles which had no cover crops grown during winter, we can define fallow sampling as maize roots grown in bulk soil. The sampling was done during the R1-RX growth stage of maize (bolting) grown in the Luvisol (01.08.2023), the Phaeozem (19.07.2023) and the Podzol (26.07.2023). The samples were extracted from the profiles using a spatula and collected in plastic zip-lock bags. Until shipment to the laboratory, samples were stored in ice coolers containing dry ice in order to preserve bacterial communities and the metabolic picture for distinct sampling time points. In the laboratory, all samples were stored at -80 °C until further processing.

### Soil physicochemical properties

Soil microbial biomass carbon (C) and nitrogen (N) were determined using the chloroform fumigation extraction method [64, 65]. In brief, 7.5 g of soil was fumigated with chloroform for 24 h and then extracted with 30 mL of 0.05 M K_2_SO_4_ on a shaker for 1 h. In parallel, 7.5 g non-fumigated soil sample was treated with the observations of the fallow samples. C and N were measured with the N/C 2100 TOC/N analyser (Analytik Jena, Jena, Germany). MBC was calculated as the difference between extracted C from fumigated and non-fumigated soil with a conversion factor (*k_C_*) of 0.45 [66]. MBN was calculated as the difference between extracted N from fumigated and non-fumigated soil with a conversion factor (*k_N_*) of 0.54 [66, 67]. The MBC and MBN were presented as µg g^-1^ dry soil. Soil density and moisture content was also measured alongside.

Soil cylinder samples were taken to measure soil bulk density at each soil depth. Soil moisture content was also measured by water content sensors (Teros 10, Meter Group, München, Germany) which were installed in the control plots (in 0-30 cm, 30‒60 cm, and 60-90 cm depth under rainout shelter and rainfall side) at each experimental site.

Once the physicochemical parameters were measured, partial Mantel’s tests and structural equation models (SEM) were applied to understand the effects of soil physicochemical properties (pH, bulk density, total organic C and total N content, soil moisture content) on the bacterial diversity in the soil using the R package *microeco* (v1.10) [68]. Additionally, the soil physicochemical proteins were correlated to individual bacterial phyla along the different soil types and soil moisture conditions to understand the impacts of these factors on the individual communities.

### DNA extraction and sequencing of 16S rRNA gene amplicons

Bacterial communities in the root-vicinity and samples were analysed by sequencing of 16S rRNA gene amplicons (2×150 bp) on an Illumina NextSeq™ 550 (Illumina, San Diego, CA, USA). DNA was extracted from 0.25 g of soil using the DNeasy^®^ PowerSoil^®^ Pro Kit (QIAGEN GmbH, Hilden, Germany). PCR amplicons of the V3 region of the bacterial 16S rRNA gene were prepared using the forward and reverse primers 341F and 518R [69] and the NEBNext^®^ Ultra™ II Q5^®^ Master Mix (New England Biolabs GmbH, Frankfurt, Germany). Sequencing libraries were prepared from 100 ng of DNA according to the Illumina protocol. Dual index adapters for the sequencing were attached using the NEBNext Multiplex Oligos for Illumina. The final concentration of the libraries was 2 nM after pooling. We sequenced triplicates of samples from each soil depth and root-vicinity source per cover crop variation plot for all three sampling sites (16S rRNA, *n*=150).

The sequencing data were analysed using QIIME2 v2023.5 [70]. First, the raw sequence reads were demultiplexed and quality-filtered (q-score ≥ 25) using the q2-demux plugin, followed by denoising with DADA2 [71] (via q2-dada2). Both the 16S forward and reverse sequences were trimmed at 150 bp. All amplicon sequence variants (ASVs) were aligned with mafft [72] (via q2-alignment), and then maximum-likelihood trees were constructed using FastTree2 [73] (via q2-phylogeny). We chose ASV-based methods over OTU approaches to limit the effect of spurious taxa on diversity indices [74]. Taxonomic assignment of bacterial ASVs was carried out using the q2-feature-classifier [75] and the classify-sklearn Naïve Bayes taxonomy classifier against the Greengenes2 “99 % OTU reference sequences” [76]. ASVs with a relative abundance of < 0.01% were defined as rare taxa. The identified bacterial phyla were categorised into r- and *K*-strategists based on literature surveys [77-80]. Phyla comprising many r- and *K*-strategists (e.g. *Pseudomonadota*,) and unassigned ASVs were categorized as ‘r- or *K*-strategist’.

### Quantitative PCR (qPCR)

The copy number of the 16S rRNA gene per gram of soil was quantified by SYBR^®^ Green-based qPCR using a 7500 Fast Real-Time PCR System (Applied Biosystems™, Thermo Fisher Scientific, Waltham, MA, USA). Aliquots of the same DNA extract utilised in amplicon sequencing were used for qPCR. Dilutions of template DNA were used to compensate for the effect of PCR inhibitors in the samples. Each sample was analysed in triplicate. PCR amplicons of the *Escherichia coli* V3 region was used as standards. Each 20 µL reaction contained 1 µL of template DNA, the forward and reverse primers 341F and 518R for 16S rRNA [81] without adapter nucleotides and Luna^®^ Universal qPCR Master Mix (NEB). Reaction conditions were an initial denaturation for 1 min at 95°C, followed by 40 cycles of denaturation at 95°C for 15 s and extension at 60°C for 30 s. The melting curve was recorded in the temperature range of 60°C to 95°C. The gene copy numbers per gram of soil were determined in comparison against the standard essentially as before [82]. The average efficiency value was 97.2 ± 5.4%.

### Metaproteomics analysis

At each timepoint, samples were collected separately from three plots for each cover crop variation at the analysed soil depths and root-vicinity sources and used for proteomic analyses following a previously described protocol [9]. Approximately 4 g of soil (from the same set of samples used for 16S rRNA sequencing) was used for protein extraction using the SDS buffered-phenol extraction method as previously described [9]. The protein extract was purified using 1-D SDS-PAGE, and then the extract was further digested with trypsin. A nano-HPLC system (UltiMate™ 3000 RSLCnano system, Thermo Fisher Scientific, Waltham, MA, USA) was used to separate the cleaved peptides. The system was connected to a Q-Exactive HF Orbitrap LC-MS/MS system (Thermo Fisher Scientific) equipped with a nano electrospray ion source, Triversa NanoMate^®^ (Advion, Ithaca, NY, USA). We linked the MS data to an in-house generated proteome database containing all the defined proteomes in UniProt for the bacteria identified in the samples by 16S rRNA amplicon sequencing. The database search was performed with Proteome Discoverer™ (v2.5.0.8, Thermo Fisher Scientific) using the SEQUEST-HT algorithm. The precursor mass tolerance of the MS was set to 10 ppm, and the fragment mass tolerance of the MS/MS was 0.02 Da. Carbamidomethylation of cysteine was considered fixed, and oxidation of methionine was set as a dynamic modification. Enzyme specificity was set to trypsin with up to two missed cleavages allowed using 10 ppm peptide ions and 0.02 Da MS/MS tolerances. Only rank-one peptides with a Percolator-estimated false discovery rate (FDR) < 1% were accepted as identified. The GhostKoala [28], KEGG [28] and COG [83] databases were used for protein functional annotation. Pathways with a minimum of two proteins and a minimum coverage of five percent were selected for downstream processing. We analysed the measured label-free quantification (LFQ) intensities of the identified proteins and determined the functional metabolic pathways to detect changes along the different sampling sites, soil moisture conditions, soil depths, and cover crop variations. During the construction of databases from UniProt for mapping the identified proteins to their respective taxonomic communities, we made sure of minimum redundancy with maximum relevancy to negate repetitive identification of already-measured proteins, which would skew the observations. In the analysis, proteins that were unambiguously identifiable by unique shared peptides and could mapped back to the UniProt reference database were used for quantifications. Using the ENTREZ key, taxonomic identities were obtained from the NCBI database [84, 85] for the identified individual proteins using unique protein identifiers called KO numbers (of KEGG). Linking the NCBI database to our datasets of identified proteins provided us with all the taxonomic information regarding the sources of the identified proteins. Each functional pathway had unique KEGG and COG identifiers, which were linked to proteins to connect the respective functional pathways and further use the NCBI-linked KO identifiers for preparing integrated datasets of taxonomic communities and functional pathways. The LFQ values were highly variable for the identified protein groups, and so they were normalised by log2-transformation using the *log* function of base R (v4.3.1) [86] prior to any graphical representations or statistical significance tests. After the categorisation of proteins into different functional pathways, we tried to incorporate our observations into the specific cycles of interest in order to understand the changes inflicted by soil property and moisture content after the introduction of re-usage of cover crop root biopores and study the microbial response. Further, to observe if environmental stress impacted the biochemical pathways post-root channel reuse, we quantified the fold-change of LFQ intensities of identified proteins under drought against the proteins from rainfall-fed conditions. Additionally, to evaluate the changes in the abundance of proteins of interest after introducing cover crop root channel re-use, differential abundances of each protein was quantified, where their values for cover crop variations (*Brassicaceae*/*Poaceae* and *Fabaceae*/*Poaceae*) were subtracted from the values from fallow. Integrating the taxonomic information with protein datasets provided an overview of the role of these communities in the biochemical pathways with changeable soil composition and environmental conditions such as drought.

### Statistical data analysis

We used R (v4.3.1) [86] to perform all statistical analyses of the sequencing and the metaproteomics data. All measures of significance were calculated using multivariate analysis of variance (ANOVA) and linear mixed models, followed by Tukey’s range post-hoc test (TukeyHSD) with package *stats* (v3.6.2) and *rstatix* (v0.7.2) [87]. In the 16S rRNA sequencing analysis, the ASV abundance tables were filtered with total-frequency-based filtering based on 95% sequence identity (via q2-feature-table summarize) and rarefied at 30,000 sequences to ensure equal sampling depth and sorting in the maximum number of samples for diversity analyses. Alpha and beta diversity metrics were calculated using the packages *phyloseq* [88], *metacoder* and *microbiome* [89] from R (v4.3.1) [86]. Shannon richness measured for each cover crop variation at different sampling sites and conditions was used for estimating alpha diversity richness. Pielou’s evenness is the most widely used diversity evenness index in the ecological literature, which also takes into estimation [90]. For beta diversity, we used Jensen-Shannon divergence (JSD) units [91] and visualised differences via Principal Coordinate Analysis (PCoA) using the *vegan* package (v2.6-4) [92]. Using multivariate anova of variance (MANOVA), we evaluated the significantly different cover crop variations using the sampling sites, sampling conditions and sampling depths as random effects and cover crop variations as the fixed effect. This was followed by Tukey’s HSD for evaluating MANOVA test outcomes with parameters of cover crop variations, depth, sampling sites, soil moisture conditions and bacterial phyla. A four-way permutational multivariate analysis of variance (PERMANOVA) among the parameters of source, variation, depth, and time points was used to quantify the significance among the parameters based on UniFrac distance using *adonis2* of *vegan* package (v2.6-4) [92]. For bacterial abundances under different parameters, the significance between the parameters was represented using the Compact Letter Display (CLD) [93] with the help of the *emmeans* package [94]. For metaproteomics, the significantly different cover crop variations or proteins of different metabolic pathways or bacterial phyla were calculated using MANOVA, using source, variation, depth, time points, and bacterial phyla as fixed factors. Upon determination, they were represented by significant stars based on the adjusted *p*-values (\**p* < 0.05, \*\**p* < 0.01, \*\*\**p* < 0.001, \*\*\*\**p* < 0.0001). All figures were generated in RStudio using the packages *ggplot2* (v3.4.2) [95], *cowplot* (v1.1.1)[96], *circlize* (v0.4.16) [97], *WeightedTreemaps* and *phyloseq* (v1.44.0) [88]. Other integrated packages used for statistical analyses and figure generation were *tidyverse* (v2.0.0), *dplyr* (v1.1.3), and *splitstackshape* (v1.4.8) [98].

## Data availability

The authors declare that the data supporting the findings of this study are available within the article, its supplementary information files, and in open access suppositories. The raw sequencing data and the respective metadata generated from this study are available under the NCBI BioProject ID PRJNA1240274, which can be accessed using the link: https://www.ncbi.nlm.nih.gov/sra/PRJNA1240274. The raw qPCR data along with the sample metadata are available on Zenodo under the DOI: https://doi.org/10.5281/zenodo.14917324. The metaproteomics datasets generated during the current study are available in the PRIDE (PRoteomics IDEntifications Database) data repository with the sample metadata, vide PRIDE dataset identifier PXD062138 (This dataset can be accessed at https://www.ebi.ac.uk/pride with the reviewer account: Username, Password: Bxr9p0XejQlb or by using reviewer access details: Project accession: PXD062138, Token: UUKPBZmgzjNb).

## Acknowledgments

The author would like to take this opportunity to thank his institution and especially the Helmholtz-Centre for Environmental Research (UFZ)-funded ProMetheus platform for metaproteomics and support. DG has also been supported by the Helmholtz Interdisciplinary Graduate School for Environmental Research (HIGRADE). We acknowledge Kathleen Eismann for her help in sample preparation for metaproteomic assessments; Habibu Aliyu, Florian Lenk and David Thiele for their assistance during Illumina NextSeq™ sequencing; Sven B. Haange for the R-scripts for analysing Proteome Discoverer outputs in metaproteomics; Stephan Schreiber for advice during NextSeq™ sample preparations; Henrik Füllgrabe, Juanjuan Ai and Tobias Stürzebecher for their assistance in fieldwork; and Matthias Bernt for his advises regarding data analysis.

## Author information

### Authors and Affiliations

Department of Molecular Toxicology, Helmholtz Centre for Environmental Research – UFZ

GmbH, Permoserstraße 15, 04318 Leipzig, Saxony, Germany

Debjyoti Ghosh, Martin von Bergen, Nico Jehmlich

Institute of Plant Nutrition and Soil Science, Department of Soil Science, Christian-Albrechts-University Kiel, Hermann-Rodewald-Straße 2, 24118 Kiel, Schleswig-Holstein, Germany Yijie Shi, Iris M. Zimmermann, Sandra Spielvogel

Institute of Crop Science and Plant Breeding, Agronomy and Crop Science, Christian-Albrechts-University Kiel, Am Botanischen Garten 1-9, 24118 Kiel, Schleswig-Holstein, Germany

Katja Holzhauser

Institute for Biochemistry, Faculty of Biosciences, Pharmacy and Psychology, University of Leipzig, Brüderstraße 34, 04103 Leipzig, Saxony, Germany

Martin von Bergen

German Centre for Integrative Biodiversity Research (iDiv) Halle-Jena-Leipzig, Puschstraße 4, 04103 Leipzig, Saxony, Germany

Martin von Bergen

Institute for Biological Interfaces, Karlsruhe Institute of Technology, Hermann-von-Helmholtz

Platz 1, 76344 Eggenstein-Leopoldshafen, Baden-Württemberg, Germany

Anne-Kristin Kaster, Jochen A. Müller

Geo-Biosphere Interactions, Department of Geosciences, University of Tübingen, Schnarrenbergstraße 94-96, 72076 Tübingen, Baden-Württemberg, Germany Michaela A. Dippold

### Contributions

Conceptualisation: NJ, JAM, MAD, SS; Methodology: NJ, JAM, IMZ, MAD, SS; Software: DG; Validation; DG, NJ, JAM; Formal analysis: DG; Investigation: DG, YS, KH; Resources: NJ, JAM, IMZ, AKK, MvB; Data curation: DG; Writing-Original Draft: DG; Writing-Review & Editing: DG, NJ, JAM, IMZ, MAD, YS; Visualisation: DG; Supervision: NJ, JAM; Project administration: IMZ; Funding acquisition: NJ.

### Correspondence

Correspondence to Nico Jehmlich and Jochen A. Müller.

## Ethical declarations

### Consent for publication

Not applicable.

### Ethics approval and consent to participate

Not applicable.

## Supplementary information

Additional figures (Fig. S1-S10) and tables (Tables S1-S12) are supplementary material related to this article. The supplementary text and figures are provided in a single .docx file and the supplementary tables as individual sheets in a single .xlsx file.

