## Supplemental_text_and_figures_S1-S10 for "Cover crop root channels promote bacterial adaptation to drought in the maize rhizosphere"

Ghosh et al.

### Materials and Methods

### Confirmatory factor analysis

We implemented a confirmatory factor analysis approach while designing the model where we aimed at testing the maximum likelihood of data fit to the hypothesised equation pathway to interpret how physicochemical properties affect bacterial presence and role in the rhizosphere. We created a latent variable for the soil quality which included pH, bulk density and soil moisture content. Regressions with the latent variable was estimated with the soil content of total organic C and total N, and the bacterial richness. The SEM model equation looks like this:

*Latent variable:*

*Soil Quality =~ pH + Density + Moisture*

*Biological Activity =~ Community Richness + Protein Richness*

*Biological Activity ~~ Soil Quality*

*Regressions:*

*Carbon ~ Density + Soil Quality*

*Nitrogen ~ Density + Soil Quality*

*Community Richness ~ Soil Quality*

The fitness of the equation model and the structural relationships with the data of the parameters were performed using the *lavaan* R package (v0.6-19)^1^. The quality of the model was further verified using comparative fit index (CFI) (0.93_CFI_ > 0.9, which is the baseline for reasonable fit), standardised root-mean-square residual (SRMR) (anything < 0.8 is a good fit indicator) and low Akaike information criterion (AIC). The *lavaanplot* R package (v0.8.1)^2^ helped us generate the graphical representation of the model, with the estimates and the significant associations.

### Supplementary Figures

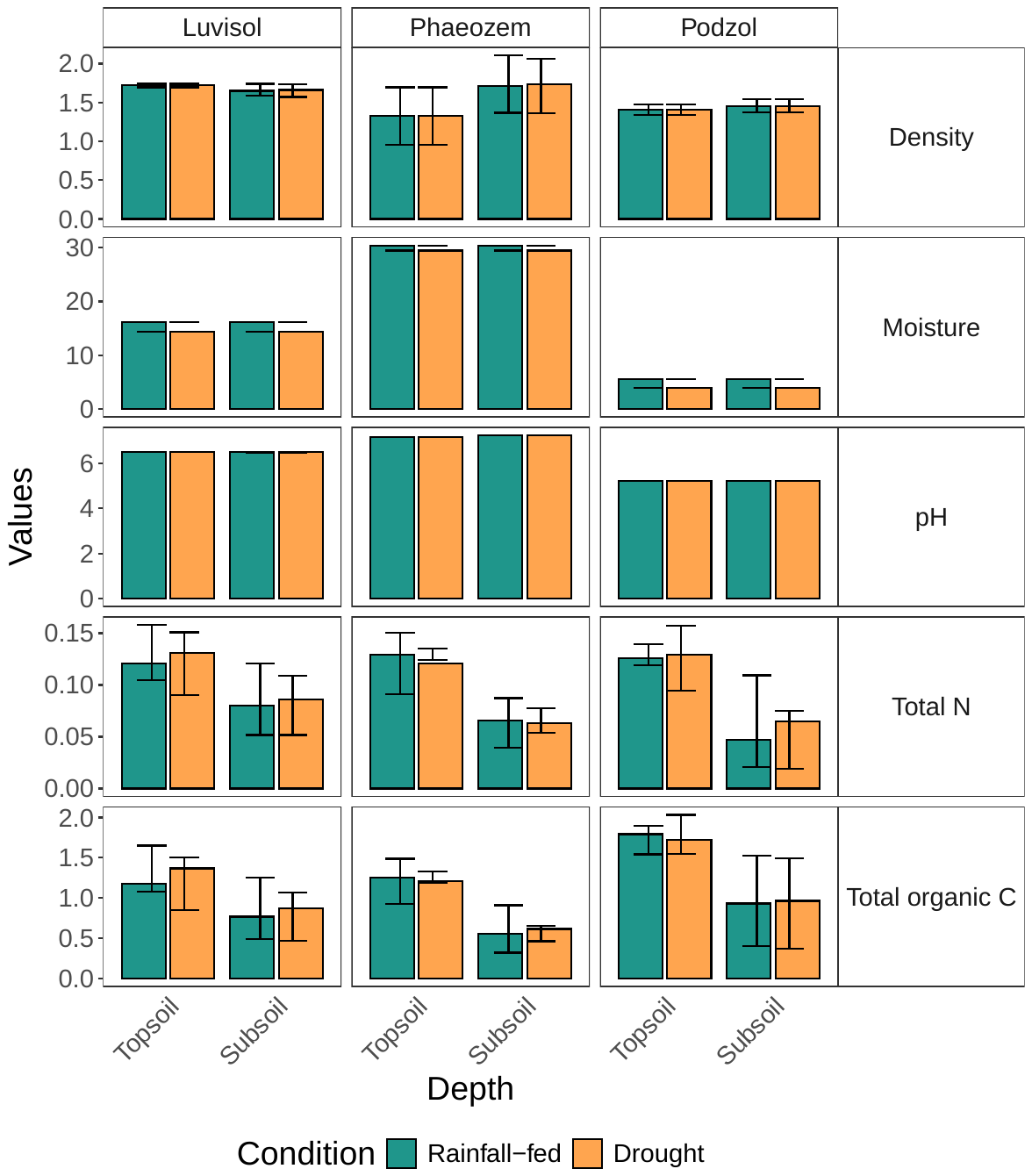

Figure S1: A bar-chart representing the different values of the physicochemical properties that have been measured at the three soil sampling sites with different soil types – Luvisol (Hohenschulen), Phaeozem (Reinshof) and Podzol (Karkendamm). All values are provided in Table S1.

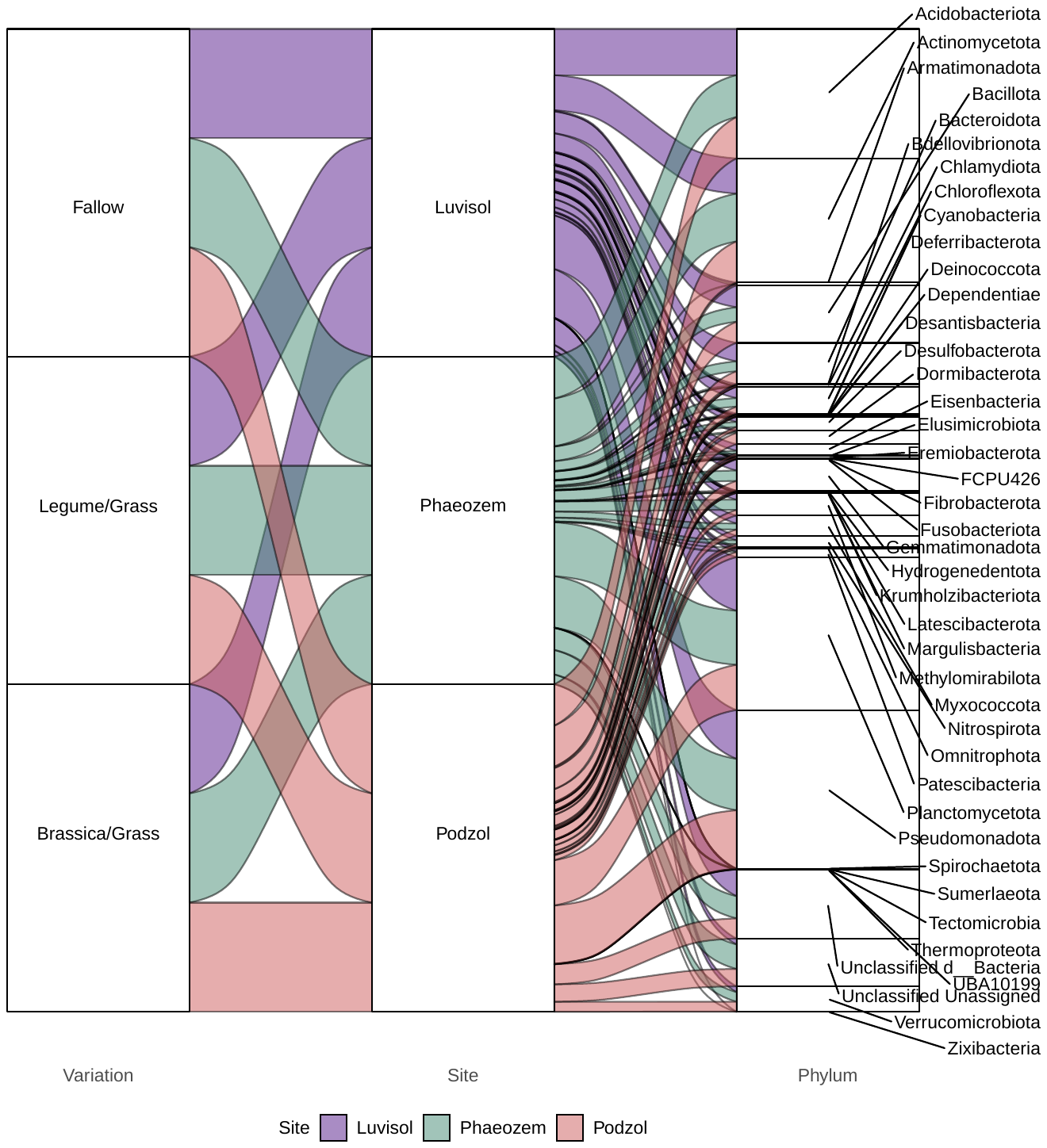

Figure S2: An alluvial flowchart representing the different variations and the identified bacterial phyla in each one of them at the three soil sampling sites.

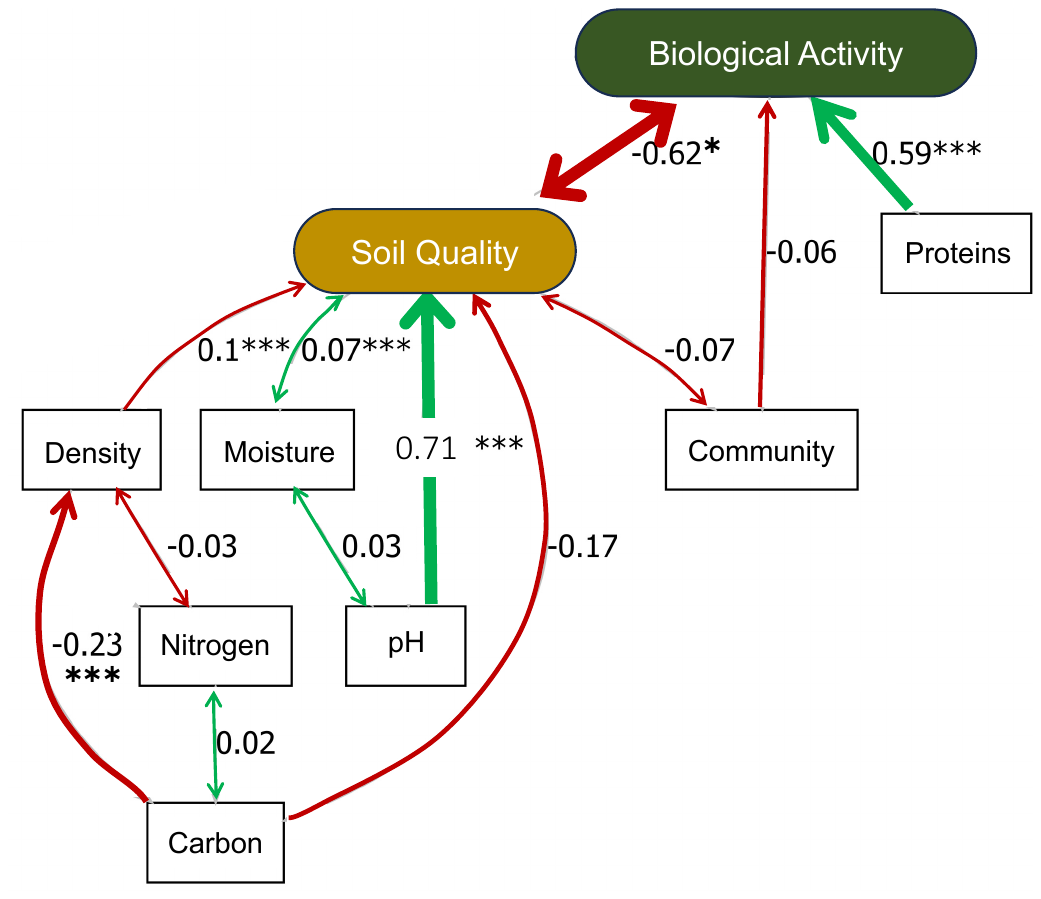

Figure S3: The conceptual confirmatory factor analysis (CFA) model evaluating the probable relationships between the soil physicochemical properties with the parameters of biological activity (bacterial and protein richness). The numbers next to the edges are estimates of the weightage of the relationships between the parameters. The quality of the model was further verified using comparative fit index (CFI) (0.93_CFI_ > 0.9, which is the baseline for reasonable fit), standardised root-mean-square residual (SRMR) (0.07 < 0.8, which is a good fit indicator) and low Akaike information criterion (AIC). The asterisks next to the arrows indicate significance of the interaction between the two parameters; **p* < 0.05, ****p* < 0.001. A summary of the analysis can be found in Table S12.

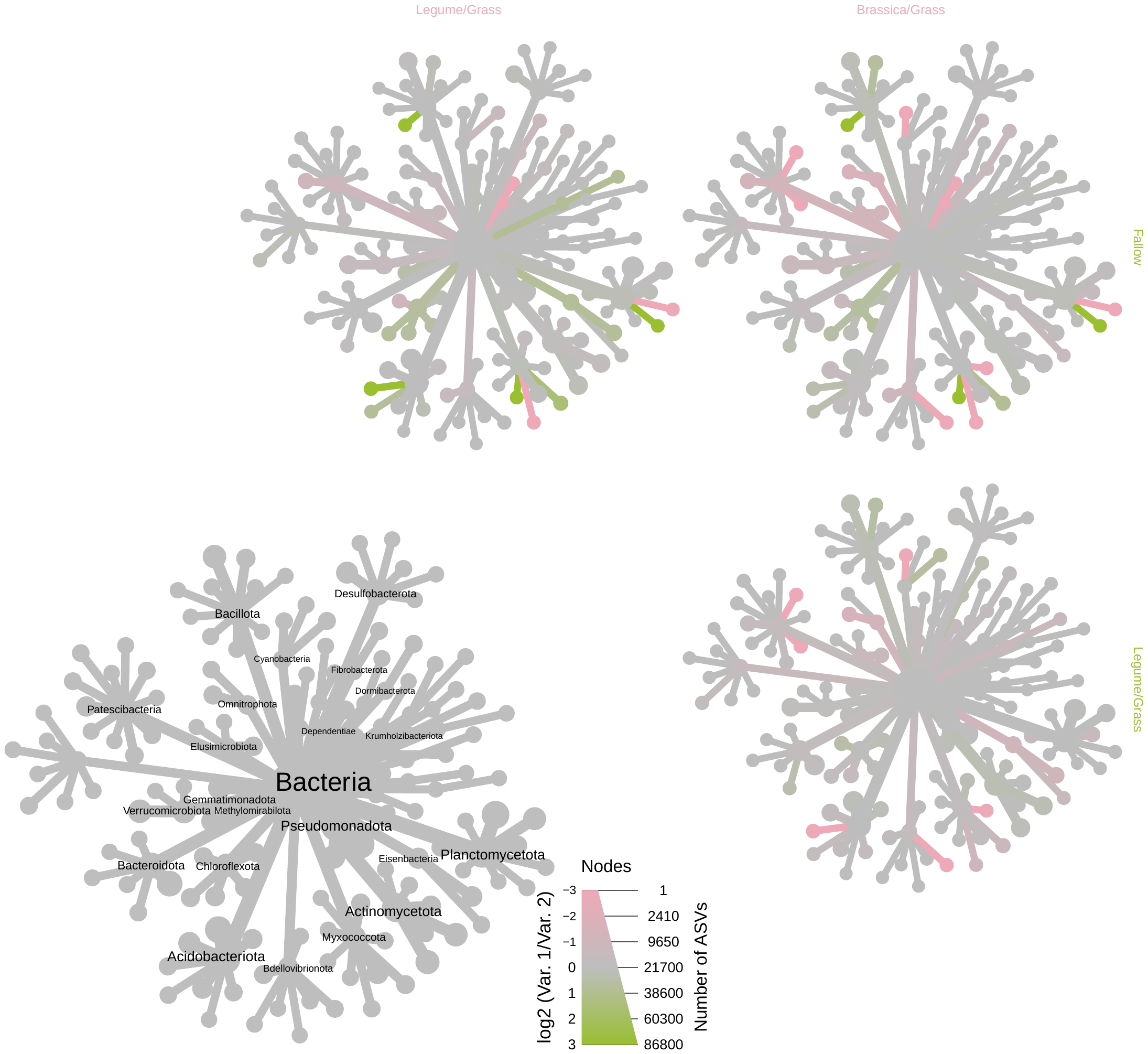

Figure S4: A heat-tree representation of the differential bacterial abundance for the cover crop variations and fallow conditions. The *log2* values of the relative bacterial abundance between treatments and fallow are shown by two colours, where ‘pink4’ corresponds to the name of the variation at the top and ‘lightgreen’ to the name on the right side (RStudio colour palettes have been used for colour representation).

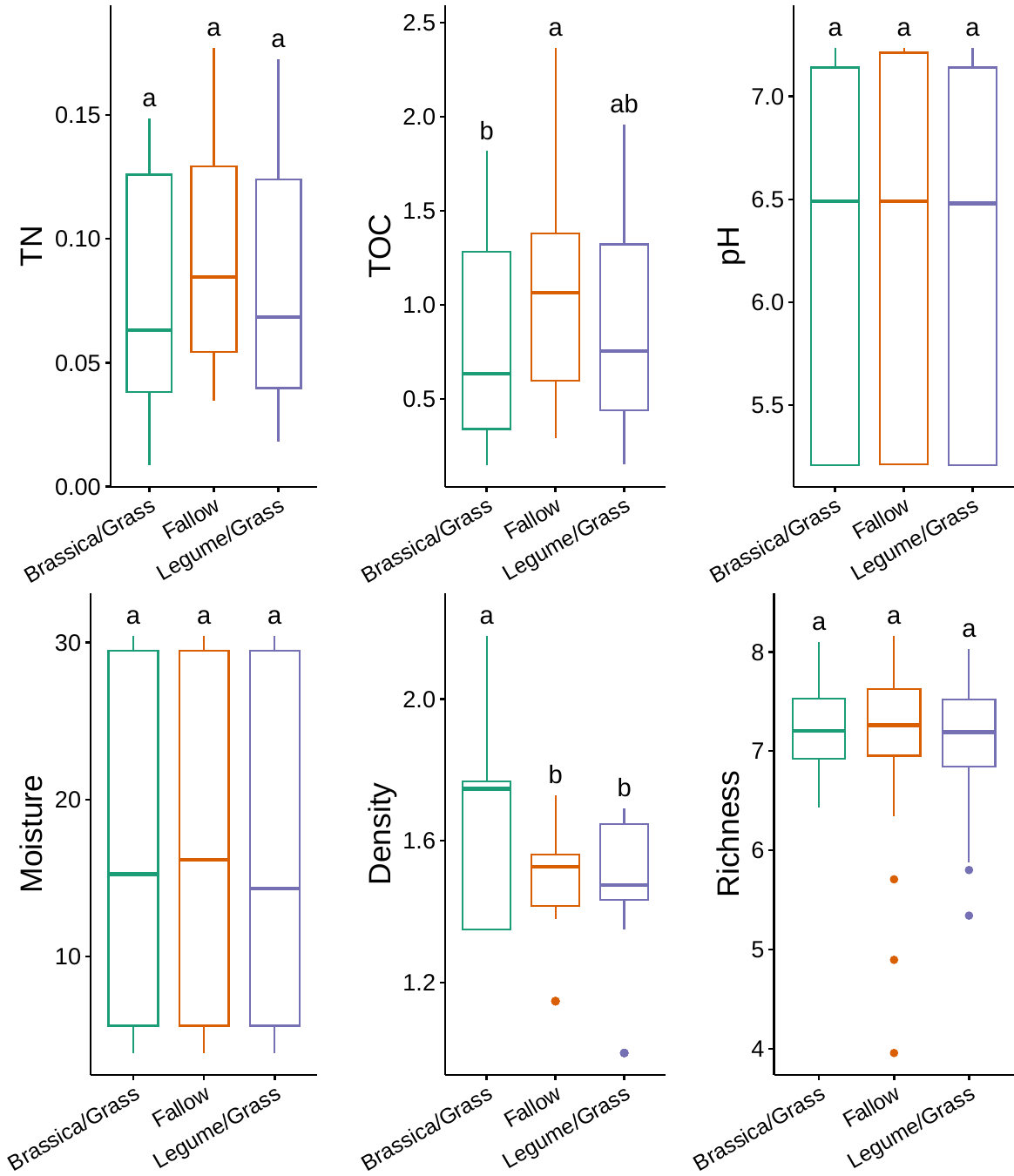

Figure S5: A faceted boxplot representation of a correlation between the soil physicochemical properties (pH, soil moisture content, bulk density, total organic carbon (TOC), and total nitrogen (TN)) plus bacterial community richness with the cover crop variations (where Fallow is the control and Legume/Grass and Brassica/Grass are treatments). Significance among the cover crop variations being used are represented by compact letter display (CLD) representations.

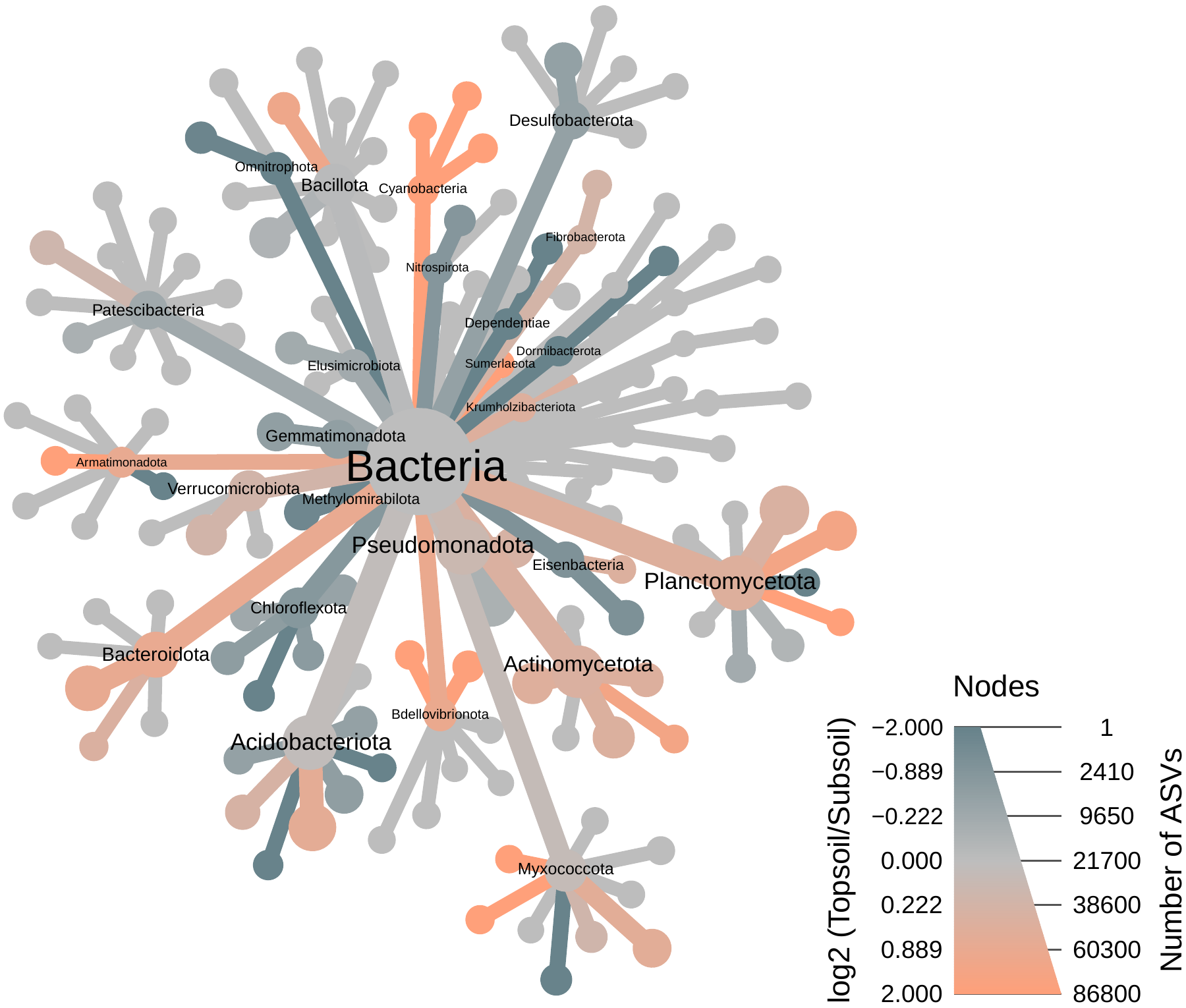

Figure S6: A heat-tree representation of the differential bacterial abundance in the two layers of soil depth – topsoil and subsoil. The *log2* values of the relative bacterial abundance between the layers are shown by two colours, where ‘lightorange’ corresponds to topsoil and ‘lightblue4’ to subsoil (RStudio colour palettes have been used for colour representation).

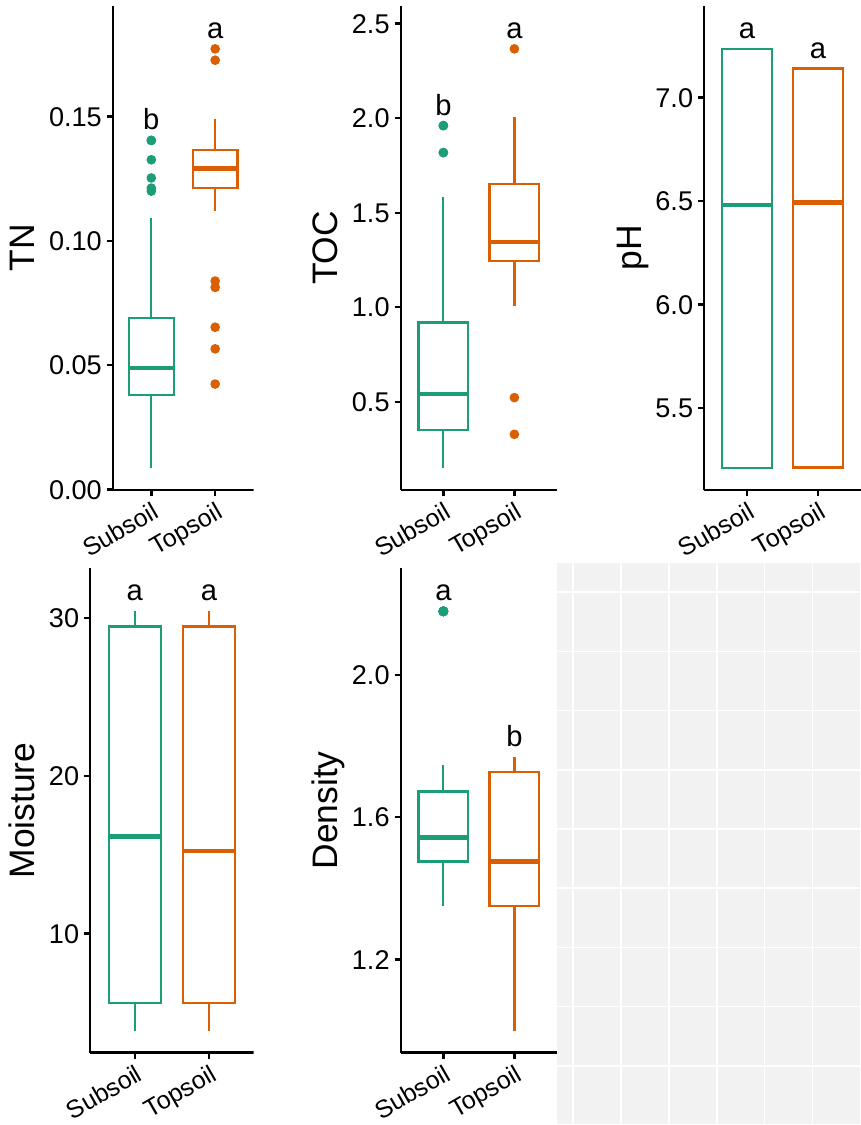

Figure S7: A faceted boxplot representation of a correlation between the soil physicochemical properties (pH, soil moisture content, bulk density, total organic carbon (TOC), and total nitrogen (TN)) plus bacterial community richness with the soil sampling depth of topsoil and subsoil. Significance among the cover crop variations being used are represented by compact letter display (CLD) representations.

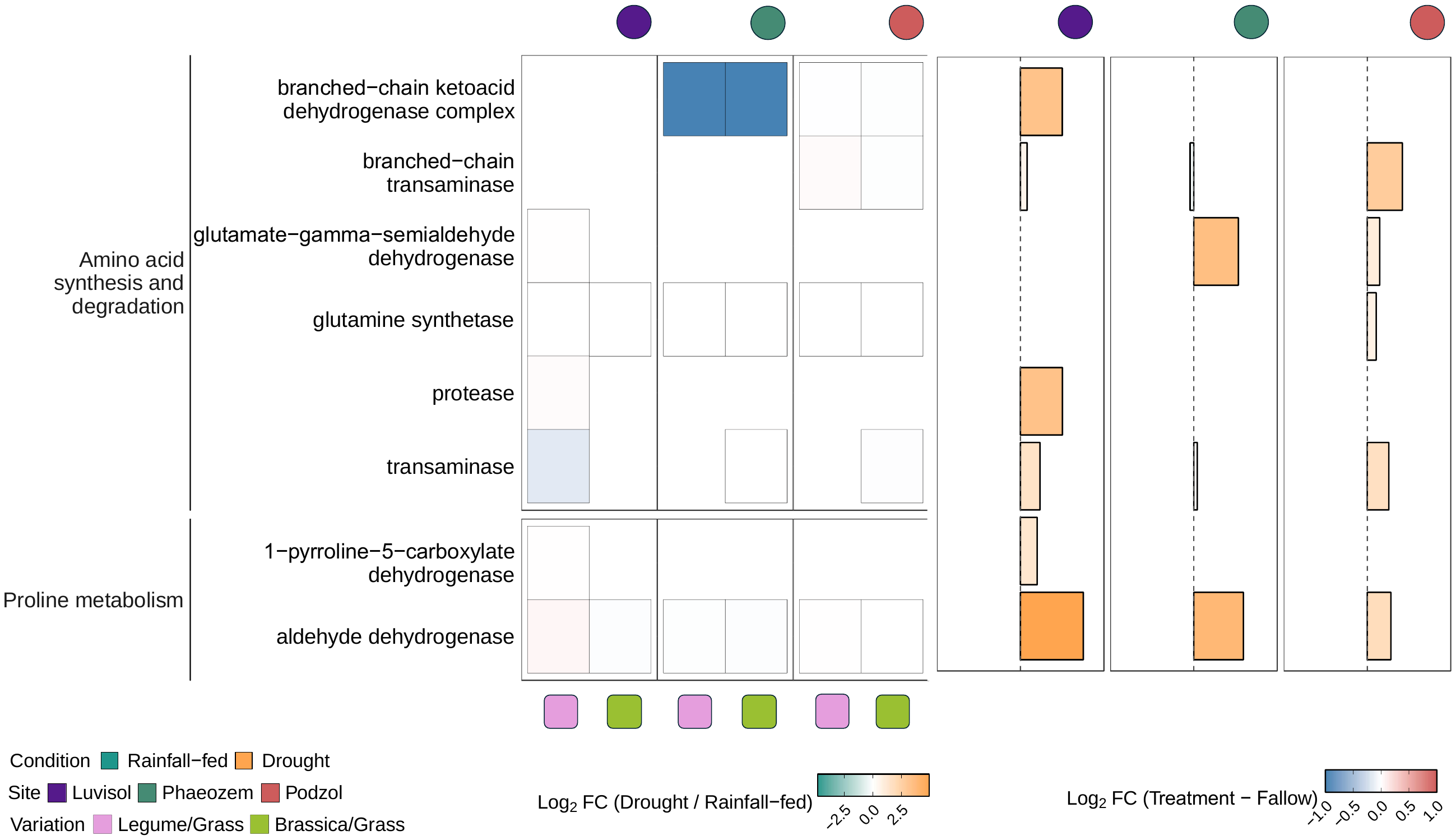

Figure S8: A summary of the expression of enzymes involved in amino acid synthesis and degradation and proline metabolism under drought and rainfall-fed conditions in our different sampling sites and the different cover crop variations.

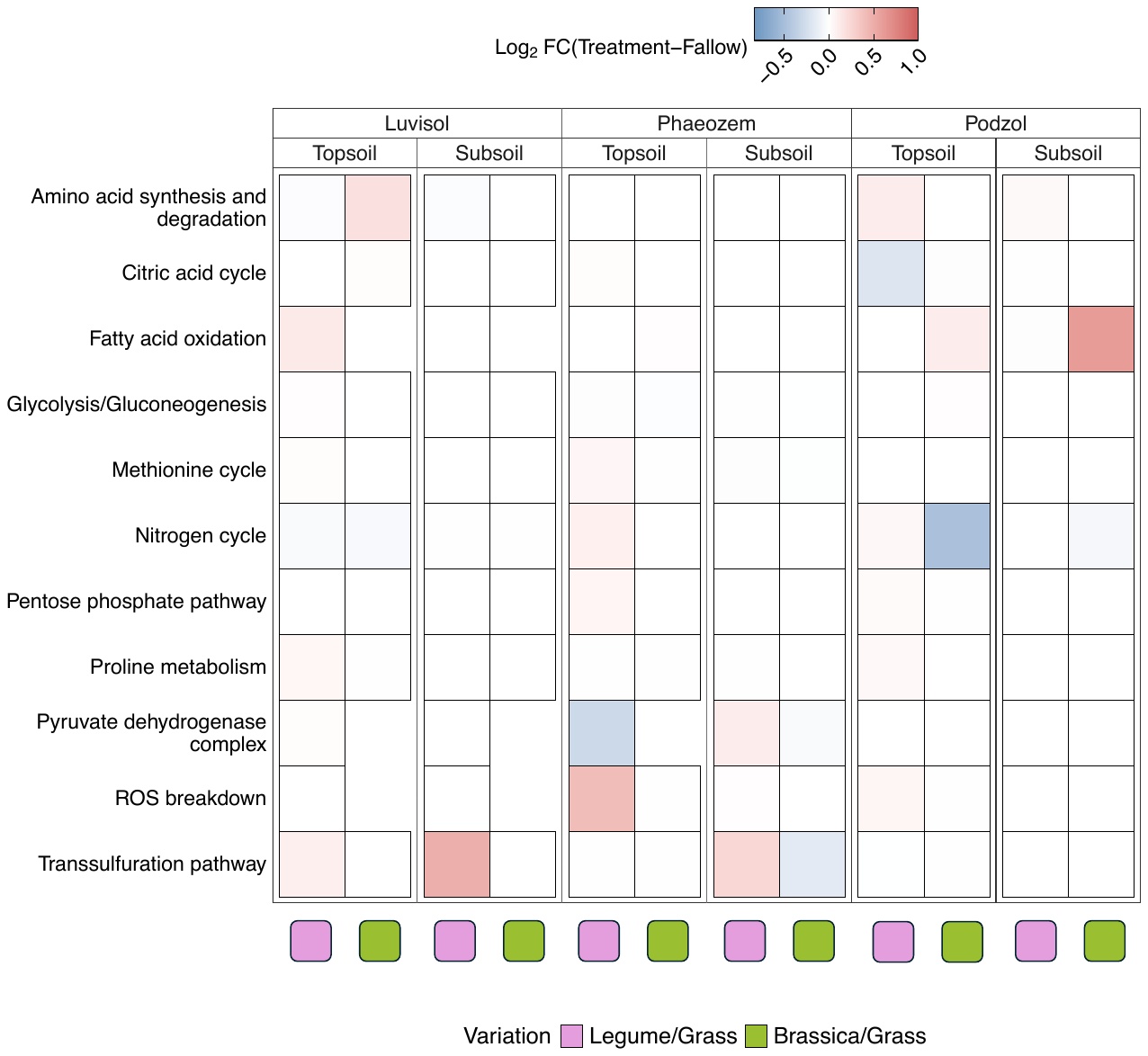

Figure S9: A summary of the differential expression of proteins of these pathways at the three sampling sites in the topsoil and the subsoil, after the re-use of the cover crop root channels. Here, the Log2FC values of the treatments (Legume/Grass and Brassica/Grass) have been deducted by the values from fallow to see the net change in the expression of proteins belonging to these pathways.

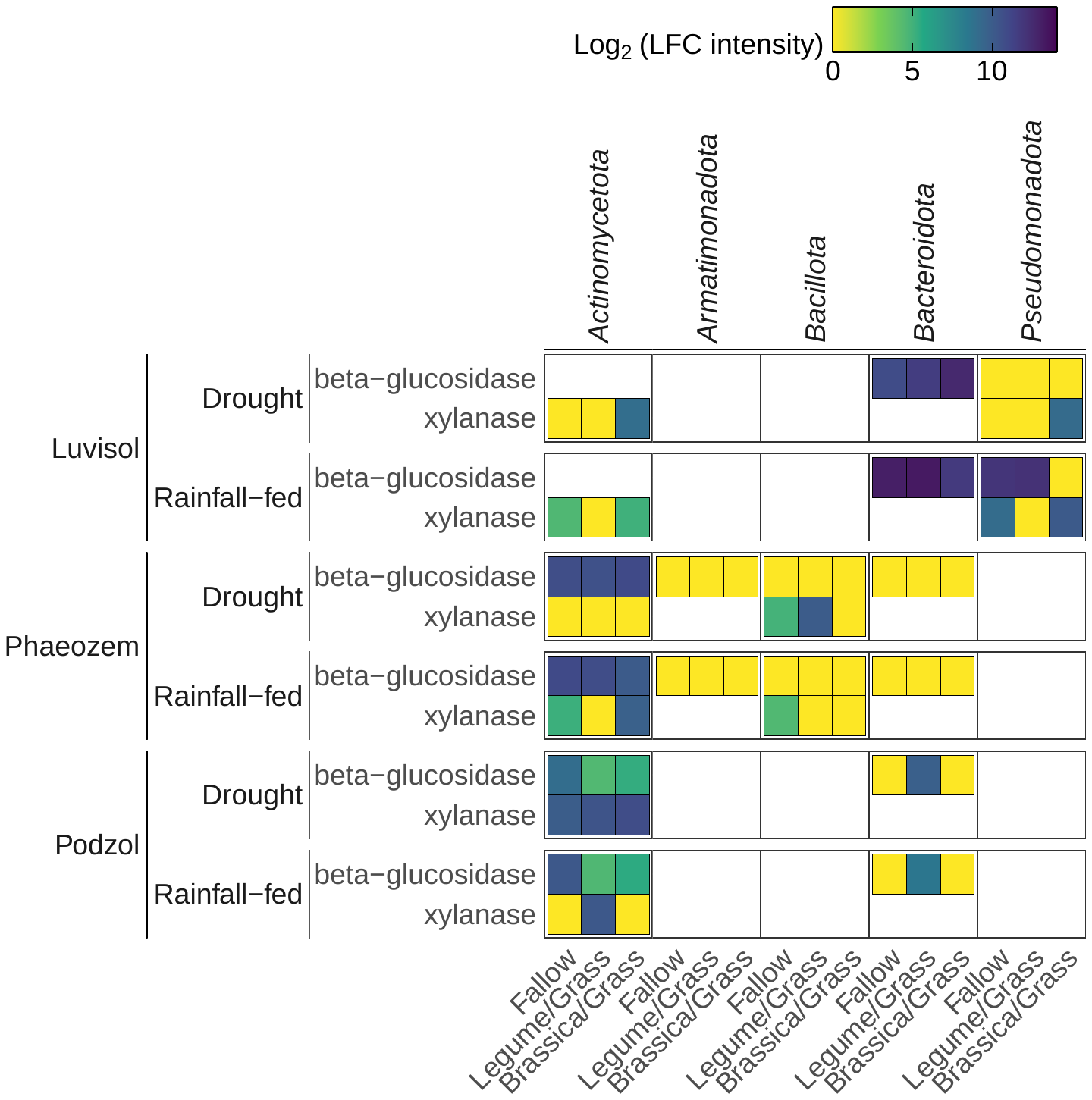

Figure S10: A summary of cellulase expression under drought and rainfall-fed conditions in our different sampling sites and the different cover crop variations. The identified cellulases were mapped to bacterial communities in order to find out which phylum plays a major role in breaking down cellulose.
