## Supplementary figures and images for "Cover crop root channels promote bacterial adaptation to drought in the maize rhizosphere"

### Supplemental_figure_S2

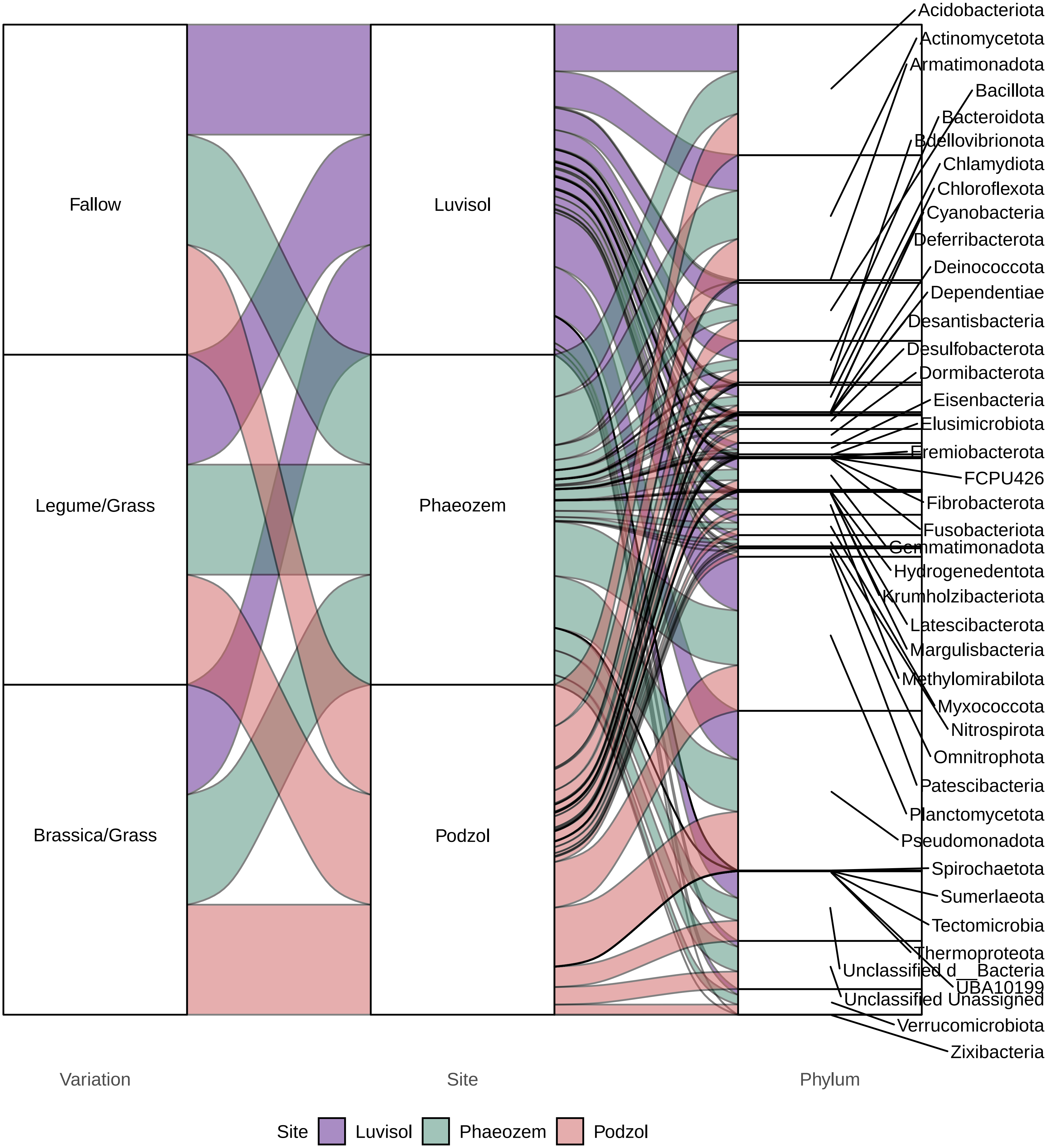
